# All-optical imaging and patterned stimulation with a one-photon endoscope

**DOI:** 10.1101/2021.12.19.473349

**Authors:** Jinyong Zhang, Ryan N. Hughes, Namsoo Kim, Isabella P. Fallon, Konstantin Bakhurin, Jiwon Kim, Francesco Paolo Ulloa Severino, Henry H. Yin

## Abstract

While in vivo calcium imaging makes it possible to record activity in defined neuronal populations with cellular resolution, optogenetics allows selective manipulation of neural activity. Recently, these two tools have been combined to stimulate and record neural activity at the same time, but current approaches often rely on two-photon microscopes that are difficult to use in freely moving animals, or one-photon fiberscopes with benchtop-based digital micromirror devices that limit system portability. To address these limitations, we have developed a new integrated system combining a one-photon endoscope and a digital micromirror device for simultaneous calcium imaging and precise optogenetic photo-stimulation (Miniscope with All-optical Patterned Stimulation and Imaging, MAPSI). Using this system, we were able to successfully image striatal neurons from either the direct pathway or the indirect pathway while simultaneously activating any neuron of choice within the field of view or synthesizing arbitrary spatio-temporal patterns of photo-stimulation in freely moving mice. We could also select neurons based on their relationship with behavior and recreate the behavior by mimicking the natural neural activity with photo-stimulation. MAPSI thus provides a powerful tool for *in vivo* interrogation of neural circuit function.

## Introduction

A major technological advance in neuroscience is the ability to record and manipulate neural activity with light. Readout of neural activity is made possible by the development of genetically encoded calcium indicators that bind to intracellular calcium and emit fluorescence that is proportional to neural activity^1^. Manipulation of neural activity can be achieved by optogenetics, using different genetically expressed light-sensitive ion channels (opsins) to activate and inactivate specific neuronal populations^2,3^. Together these techniques have revolutionized neuroscience, allowing investigators to record and manipulate neural activity with cell type specificity in behaving animals^4-7^. More recently, calcium imaging and optogenetics have been combined in 2-photon (2P) microscopy to manipulate and image neural activity at the same time ^8-12^. With this ‘all-optical’ approach, the same neurons can be recorded and stimulated simultaneously, and light delivery can be restricted to individual neurons. It is also possible to mimic normal neural activity by replaying the activity from calcium imaging using more physiologically realistic stimulation parameters ^13-16^.

Currently 2P microscopy is the gold standard for all-optical stimulation and recording at cellular resolution ^9,13,17^. However, traditional 2P microscopy has key limitations such as high cost and lack of portability, making it difficult to use in freely moving animals. Although recent work has developed a portable 2P system, it is still not possible to combine patterned stimulation and calcium imaging simultaneously^18^. While 1P patterned stimulation and imaging system have also been developed ^19-22^, they also have significant limitations, such as the need for a benchtop confocal microscope and digital micromirror devices that limit system portability.

To overcome these limitations, we developed a new 1P system for simultaneous stimulation and calcium imaging: Miniscope with All-optical Patterned Stimulation and Imaging (MAPSI). MAPSI integrates pattern stimulation and calcium recording in a single system with a small body (25 mm x 15 mm x 15 mm). It makes it possible to mimic natural physiological patterns just recorded, instead of delivering artificial stimulation patterns to all neurons expressing opsins indiscriminately. Neural activity can be recorded and analyzed in real-time while selecting and stimulating neurons that are behaviorally relevant. MAPSI thus makes it possible to perform closed loop experiments in which neural activity is maintained or shaped online.

## Results

### Design of MAPSI

The calcium imaging component in MAPSI is based on the UCLA Miniscope, which allows imaging of many neurons in freely moving animals ^23^. To integrate LED excitation of calcium indicators and laser stimulation, we modified the original Miniscope design to incorporate a DMD and an additional laser light source. DMDs can create arbitrary light patterns with extremely high spatial and temporal resolution. They have been used for precisely photo-stimulation *in vitro* ^24,25^, and with a fiberscope in freely behaving mice ^22,26^.

To record and stimulate simultaneously, we used two light sources: the first is a lime LED (540 nm-580 nm filter) for calcium sensor excitation. This LED is controlled by a constant current source capable of producing up to 200 mA with a step size of 4 mA, with up to 12 mW/mm^2^ measured beneath the gradient-index (GRIN) lens (**Supplementary Figure 1**). The second light source is an external laser (Opto Engine LLC, PSU-H-LED) that generates blue light for optogenetic excitation (473 nm). By passing the excitation light through the DMD ^25,27^, we could generate a patterned light beam, which is merged with the excitation light for calcium imaging in the main excitation path (**Figures 1A-B**). The DMD is controlled by a display controller (DLP3430) that can connect to any computer through an HDMI interface board (**Supplementary Figure 1**). The orientation of the micromirrors can be precisely controlled by a computer to create different stimulation patterns and sequences. It can be programmed to split the collimated light from the laser into individual beams, allowing the experimenter to select and stimulate extremely small areas (**Figures 2A-B & Supplementary Figure 2**). In the DMD off-state, an absorber is used to avoid light dispersion into the excitation light path.

**Figure 1.**
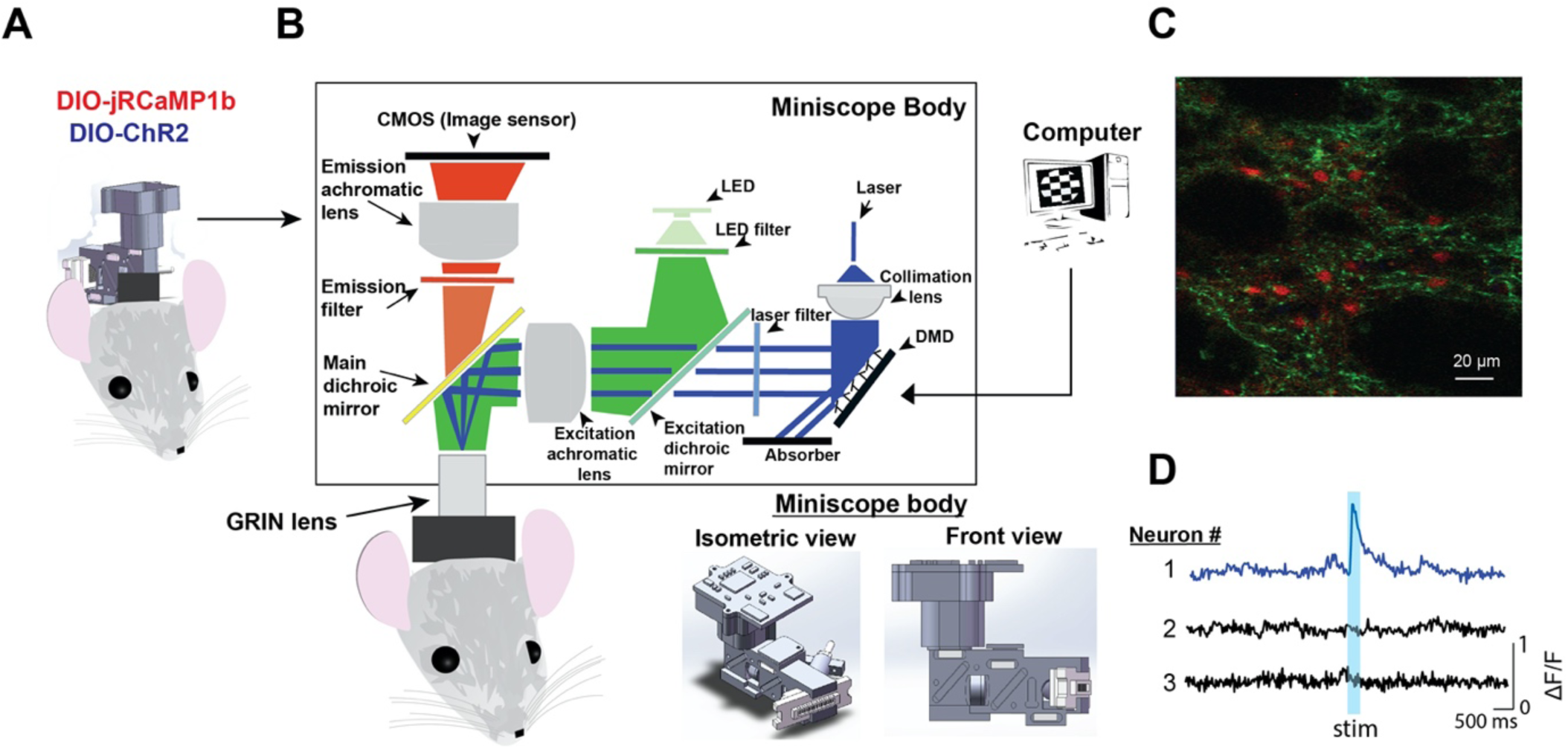
Miniscope with All-optical Patterned Stimulation and Imaging (MAPSI) **A)** Drawing of MAPSI on a mouse head. jRCaMP1b, a red shifted calcium indicator, was used for imaging, and ChR2 for optogenetics. **B)** Schematic illustration of MAPSI components. Two light sources are in the excitation path: the lime LED excites jRCaMP1b and the blue laser light excites ChR2. Micromirrors on the DMD reflect a collimated laser beam, generating arbitrary stimulation patterns created and controlled by a computer. The focal plane of the excitation achromatic lens matches the GRIN lens focal plane to ensure high resolution. The emission path records calcium activity with a CMOS sensor. **C)** Histological image showing dSPNs (D1-cre mouse) that co-express jRCaMP1b and ChR2. **D)** Calcium transients showing excitation of stimulated neuron #1, but not of nearby neighboring neurons #2 and #3.

**Figure 2.**
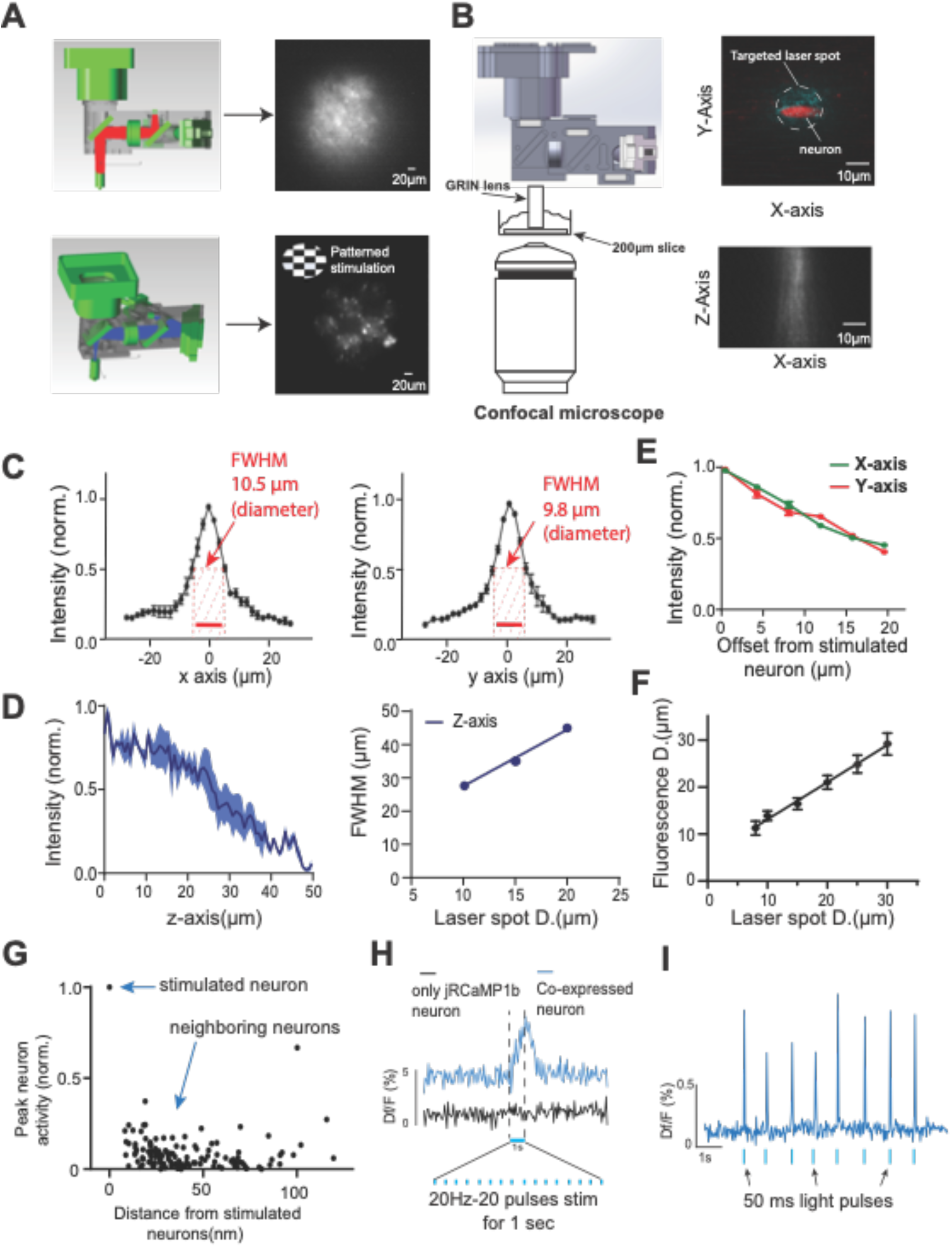
High resolution of MAPSI allows patterned stimulation and calcium imaging. **A)** Top: Schematic of the LED excitation path and the FOV (∼250 µm). Bottom: Schematic of the patterned optogenetic excitation path. A checkered illumination pattern was used to demonstrate its ability to create patterns. **B**) Left: Schematic of the setup to validate axial (z-axis) resolution. Right: The resolution of stimulated beamlets on the x-y plane and x-z plane using the confocal microscope. **C)** Lateral (x and y) resolution. Left and middle: The FWHM of fluorescence (N = 6 mice) is less than 10.5 µm on the x axis and 9.8 µm on the y axis using a 10 µm diameter spot. **D)** Fluorescence intensity decreases as a function of laser spot offset. **E**) Axial (z-axis) resolution. Left: The light intensity along the z-axis using a 10 µm diameter laser beamlet (power =70 mW/mm^2^, n = 6 stimulated neurons). Right, axial resolution (FWHM) of the photoactivation beam linearly increases with the spot diameter. **F)** There is a linear relationship between fluorescence diameter and the laser beam diameter (D). The minimum fluorescence diameter is 10 µm. **G)** Peak fluorescence of all recorded neurons as a function of distance from the stimulation location (0-500 ms after stimulation onset, 5 mice, 132 neurons, power = 70 mW/mm ^2^,) **H)** Comparison of signals from a SPN co-expressing both calcium indicator and opsin and a SPN expressing only the calcium indicator. **I)** A representative stimulated neuron showing high fidelity responses to stimulation. All error bars indicate SEM.

To obtain high resolution, the key challenge is to produce almost perfectly collimated light (< 1% deviation). We were able to achieve this by simulating its design in optic tracing software (**Supplementary Figure 3**). In the main excitation light path, the focal planes of the achromatic lens and the GRIN lens are matched so that the light will precisely target specific regions beneath the GRIN lens. With this collimated beam, photo-stimulation can maintain its precise beamlets with high resolution even after traversing long distances (20 mm) through the excitation path and the GRIN lens.

Because MAPSI is heavier than a conventional miniscope (∼7.8 g compared to ∼4 g), to help mice to carry it, we developed a commutator with a pulley to reduce the weight carried by 4 g, thus allowing free movement for long periods (**Video S1**). A direct comparison of the original UCLA Miniscope and MAPSI in the same mice moving in an open field arena is shown in **Supplementary Figure 4**. In the future, the weight can be further reduced by using a wireless DMD connector and a smaller CMOS.

### Functional capabilities of MAPSI

To test the ability of MAPSI to simultaneously stimulate and record neurons in freely moving animals, we injected both viral vectors containing Cre-dependent jRCaMP1b and ChR2-eYFP into spiny projection neurons (SPNs) in the dorsolateral striatum (DLS) ^2,8,28^. jRCaMP1b can be excited by a lime LED with a 540-580 nm excitation filter, and ChR2 can be activated using an excitation wavelength of 473 nm with a 450-490 nm emission filter from a separate laser generator. Using Cre-dependent viral vectors (AAV1-CAG-Flex-jRCaMP1b and AAV5-EF1aa-DIO-hChR2-eYFP) in D1-Cre and A2A-Cre mice, we expressed jRCaMP1b and ChR2-eYFP in direct (D1+) and indirect (A2A+) pathway neurons^29^.

MAPSI was first tested in anesthetized mice to validate that the stimulation area was similar to the calcium imaging region (**Figure 2A**). A baseplate is used to fix MAPSI on the mouse head. More neurons can be recorded and stimulated by re-baseplating to cover another region beneath the GRIN lens. Using a 1.8 mm GRIN lens, the recording FOV is circular (diameter ∼ 250 µm), allowing simultaneous calcium imaging and stimulation. The power density is 2 mW/mm^2^ for imaging excitation, and 60-80 mW/mm^2^ for optogenetic stimulation. The power density for optogenetic stimulation is similar to what is commonly used in the field (usually 40-200 mW/mm^2^) ^30,31^. It has been shown that unacceptable tissue heating is possible with prolonged stimulation at a higher power of >100 mW/mm^2^ (>3 mW light from a fiber with 200 µm diameter)^31^. In MAPSI, with a spot 10 µm in diameter for single neuron stimulation, the power is ∼60-80 mW/mm^2^, which is unlikely to create significant tissue heating.

### Axial and lateral resolution

To determine the axial (z-axis) resolution of MAPSI, we first recorded fluorescence signals from a 200 µm brain slice infected with ChR2 and RCaMP1jb using a confocal microscope, while simultaneously stimulating neurons using MAPSI. The axial resolution and depth of penetration are shown in **Figure 2B**). We found that, the full-width half-maximum (FWHM) of the stimulation beamlets is approximately 30 µm. When the laser spot is 10 µm in diameter, the fluorescence detected was almost circular with 10.5 µm FWHM on the x-axis and 9.8 µm FWHM on the y-axis (**Figure 2C**). The fluorescence linearly increased as the diameter of the illuminated area increased, and intensity decreased as the beamlets penetrated deeper into the tissue (**Figure 2D**).

To identify which neurons co-expressed both ChR2 and jRCaMP1b, the jRCaMP1b was continuously excited at low power (∼1 mW/mm^2^) to achieve a stable baseline, and ChR2 was excited at 20 Hz (20 ms pulse duration, 20 pulses, power density ∼50 mW/mm^2^). During stimulation, we were able to measure a significant increase in jRCaMP1b fluorescence signal by targeting the neuron using a 10 µm diameter spot (**Figure 2E**, 31 neurons in 6 mice. All traces are shown in **Supplementary Figure 9**). In the absence of ChR2 expression, no fluorescence change was detected (**Figure 2E**; 21 cells from 4 control mice with only jRCaMP1b expression).

### Testing in freely moving mice

MAPSI makes it possible to select individual neurons that are active during behaviors of experimental interest and play back their activity. In order to mimic naturally occurring neural activity patterns, we identified neurons that were active during a behavior of interest. If they co-expressed both ChR2 and jRCaMP1, we could then optogenetically stimulate these neurons while recording their activity (**Supplementary Figure 4C**).

We used Cre-dependent viral vectors and Cre driver lines to target either the direct pathway(striatonigral, D1-cre) or the indirect pathway (striatopallidal, A2A-cre) ^32-34^. These two well-established pathways are known to have opposite effects on basal ganglia output and behavior ^35-38^. In particular, stimulation of direct pathway spiny projection neurons (dSPNs) can produce contraversive turning behavior (away from the side of stimulation, toward the contralateral side), and stimulation of indirect pathway spiny projection neurons (iSPNs) can produce ipsiversive turning behavior (toward the side of stimulation)^35^. Thus, optogenetic manipulation of these pathways provide convenient behavioral readouts to test the efficacy of our all-optical stimulation/imaging system.

We tested MAPSI in D1-Cre mice or A2A-Cre mice, with a chronically implanted GRIN lens (1.8 mm diameter, 4.3 mm length) just above the injection site during freely-moving behavior. We generated multiple 10 µm beamlets (5 beamlets at 80 mW/mm^2^ for dSPNs, 4 beamlets at 60 mW/mm^2^ for iSPNs) for stimulation while recording calcium activity from all neurons in the FOV. Individual dSPNs or iSPNs could be robustly and selectively activated without activating neighboring neurons (**Figures 3B-C, Supplementary Figures 5-6**). Occasionally a non-stimulated neuron (e.g. neuron 6 in Figure 3C) located close to a stimulated neuron (neuron 1 in **Figure 3C**) responded almost systematically but with a longer latency (**Figure 3D**). This suggests that some neurons can be activated indirectly by the stimulation via circuit connections.

**Figure 3.**
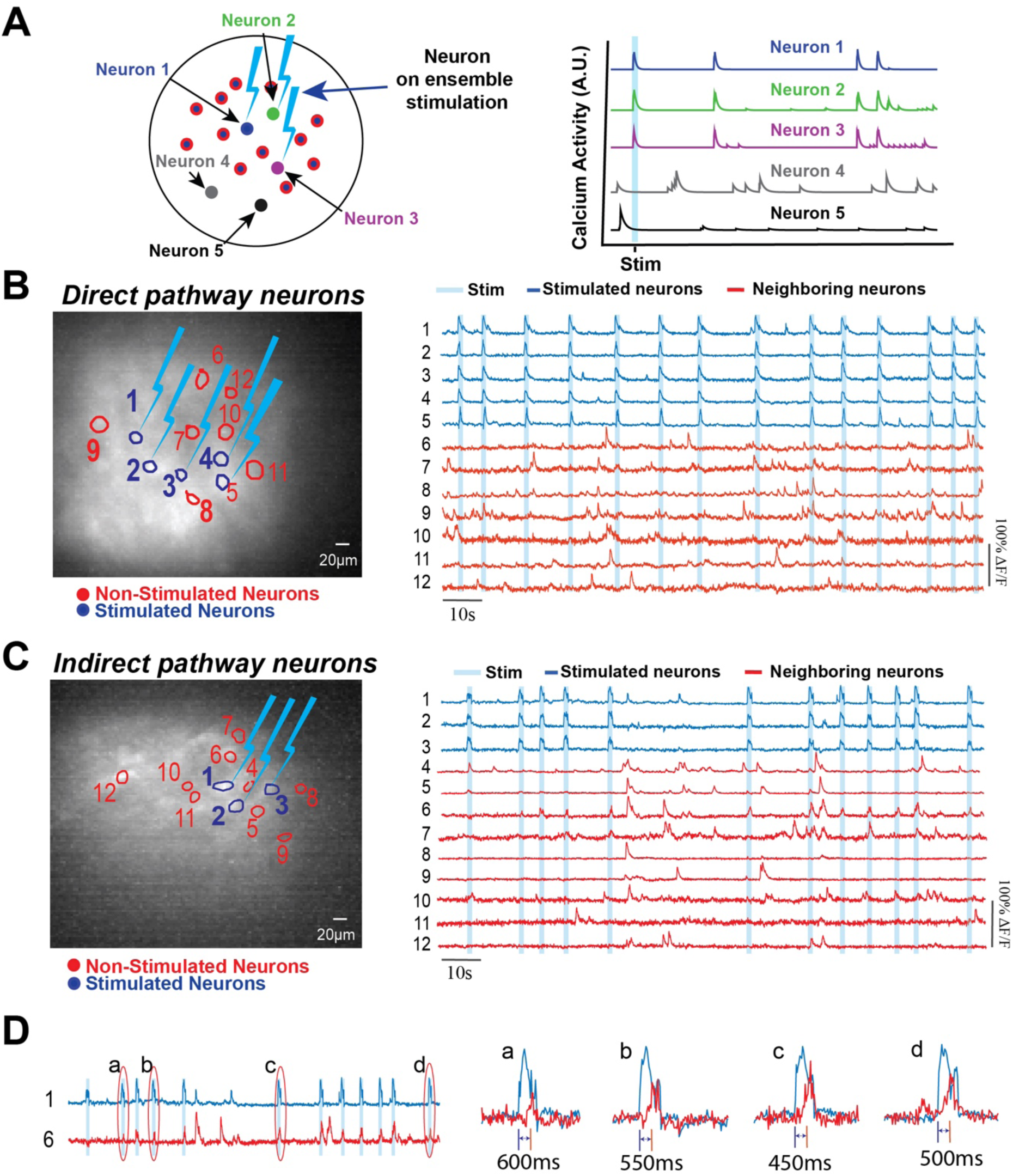
Resolution of MAPSI stimulation in freely behaving animals. **A)** Multiple neurons are selected for simultaneous stimulation. Five representative traces are shown on the right: top 3 are directly stimulated neurons, and bottom 2 are neighboring neurons that were not stimulated. **B)** Calcium signal from dSPNs expressing jRCaMP1b and ChR2. When 5 selected neurons were stimulated, neighboring neurons were not activated. **C)** Calcium signal from iSPNs co-expressing jRCaMP1b and ChR2 in a representative mouse. When stimulating 3 neurons, neighboring neurons were not excited by the stimulation. D) Comparison of neuron 1 and neuron 6 from C, showing several representative trials. Neuron 1 is directly activated by photo-stimulation. Although the neighboring neuron 6 is often activated by the stimulation, the evoked activity is highly variable, with a long latency (∼500 ms). This pattern suggests that neuron 6 is indirectly activated, presumably via some circuit connection presumably involving multiple synapses.

### Isolating behaviorally active neurons and manipulating their activity

To examine the effects of direct pathway activation, we injected Cre-dependent jRCaMP1b and ChR2-eYFP into D1-Cre mice (N = 3 mice) and only jRCaMP1b in control mice (N = 2 mice) while simultaneously recording their behavior in an open field arena (**Figure 4A**). Because D1-neurons are known to increase their activity during contraversive movements ^36,37,39^, we classified neurons that increased firing within 500 ms from the start of contraversive turning movement as turning-related neurons (28 out of 86 recorded neurons from 3 mice, 6-22 trials per mouse). The turning angle was computed after labeling each frame in DeepLabCut^40^(**Figure 4B-C**). First, we used DeepLabCut to mark two points, one on the head and one on the back of the mouse, and used these points to generate a ‘head-body’ vector. We compared 2 successive frames (50 FPS) to calculate the change in vector angle, which is used as a measure of turning.

**Figure 4.**
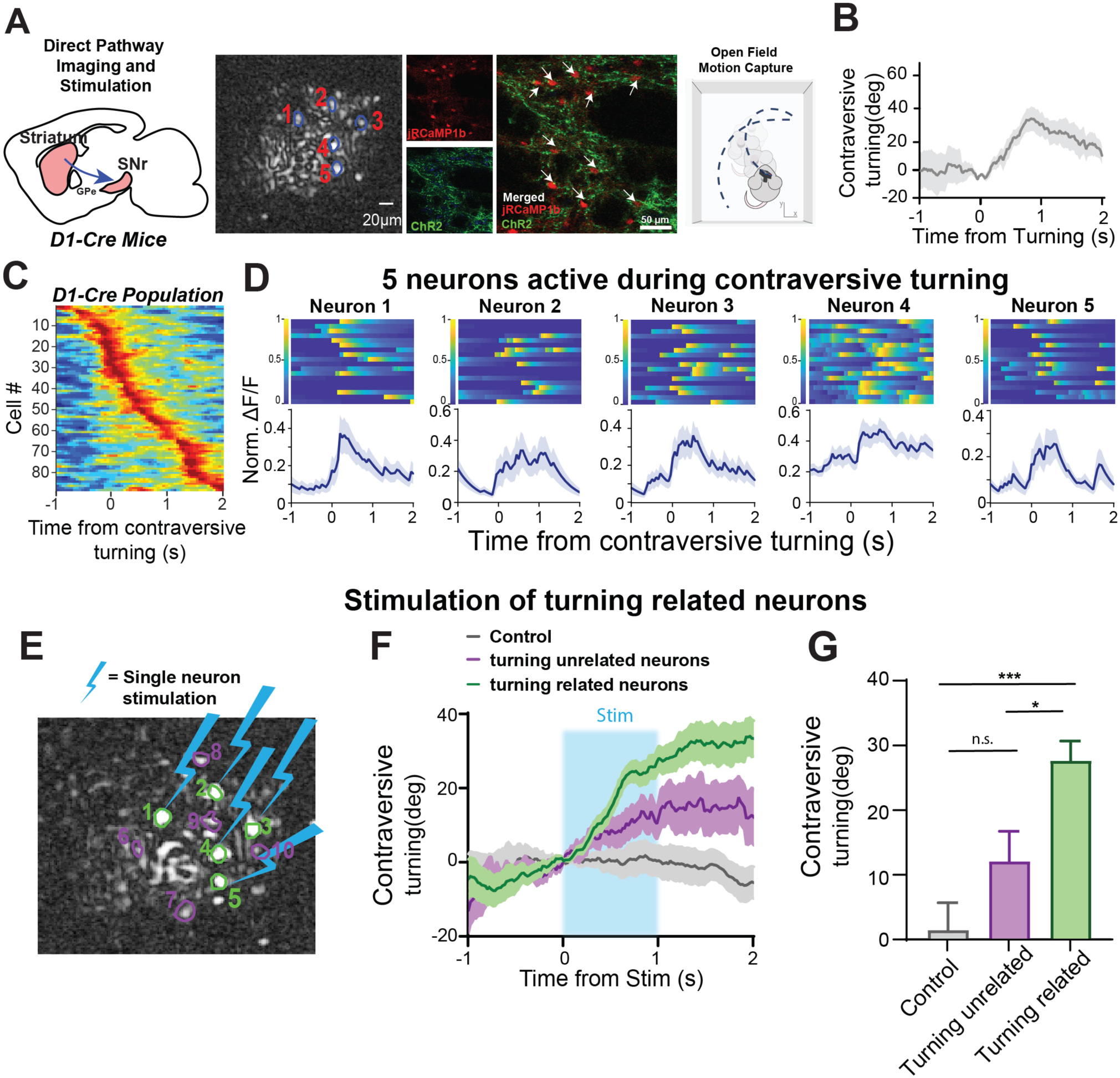
Stimulation of direct pathway neurons that are active during contraversive turning reproduces contraversive turning. **A)** 5 selected D1+ dSPNs neurons from a representative mouse in an open field arena were stimulated. These neurons co-express ChR2 and jRCaMP1b. **B)** Behavioral data was aligned to start of contraversive turning (3 D1-cre mice, 6-22 trials per mouse). **C)** All recorded dSPNs (3 D1-cre mice, 86 neurons) aligned to contraversive turning and sorted according to turning onset. **D**) Calcium activity of 5 neurons in a representative mouse activated during contraversive turning onset. **E)** The same 5 neurons were stimulated, while 5 neighboring neurons not related to turning were also stimulated. **F)** Stimulating 5 selected neurons produced more contraversive turning than stimulating unrelated neurons or stimulation in control mice (3 D1-cre mice, 5-25 trials of stimulated neurons, 4-20 trials per mouse; 2 control mice, 6-16 trials per mouse). **G)** Stimulation of 5 turning related neurons significantly increased contraversive turning angle compared to controls and unrelated neurons stimulation (one-way ANOVA: Stim groups, F(2, 50) = 9.303 p = 0.0004. Tukey’s post hoc analysis revealed stimulating contraversive turning-related neurons resulted in more turning than stimulating neighboring neurons (p = 0.0120) and controls (p = 0.0007). Stimulating unrelated neurons did not produce more turning than controls (p=0.2819). All error bars indicate SEM.

We used the derivative of the headbody vector as a measure of the angular deviation of body posture, regardless of whether the mouse was walking or not. We selected 15 of the turning-related neurons (5 neurons in each mouse) for stimulation (representative mouse shown in **Figures 4D**). Selective stimulation of these neurons elicited turning that was comparable to the mice’s natural turning (**Figures 4G & H, compare to 4B & E**). Surprisingly, stimulating 5 neurons was sufficient to produce a large turning effect. Different parameters of stimulation can also produce contraversive turning (**Supplementary Figure 7B**). The magnitude of the behavioral effect depended on the number of neurons stimulated (**Supplementary Figure 5**). In contrast, stimulating 5 neighboring neurons that are not related to turning did not produce significant turning (**Figure 4G-H**, D1-cre mice, 25 trials for stimulated neurons, 20 trials for neighboring neurons; 16 trials in 2 control mice).

We next performed the same experiment with A2A-Cre mice (N = 3 jRCaMP1b and ChR2 mice, N = 2 controls with only jRCaMP1b) (**Figure 5 A**). Activation of the indirect pathway neurons is known to produce ipsiversive turning ^35^. Out of 354 recorded iSPNs, 36 were related to ipsiversive turning. We selected 12 neurons (4 neurons in the FOV in each mouse) that showed robust excitation during ipsiversive turning (N =3 mice; **Figures 5B-C**). We then excited these turning-related neurons (n = 12 neurons from 3 mice), which produced ipsiversive turning (**Figures 5E-G**). Different parameters of stimulation can also produce ipsiversive turning (**Supplementary Figure 7**). The magnitude of the behavioral effect depended on the number of neurons stimulated (**Supplementary Figure 6**). Similar to our experiments with D1-Cre mice, we also stimulated neurons that were not active during turning, which did not produce significant ipsiversive turning.

**Figure 5.**
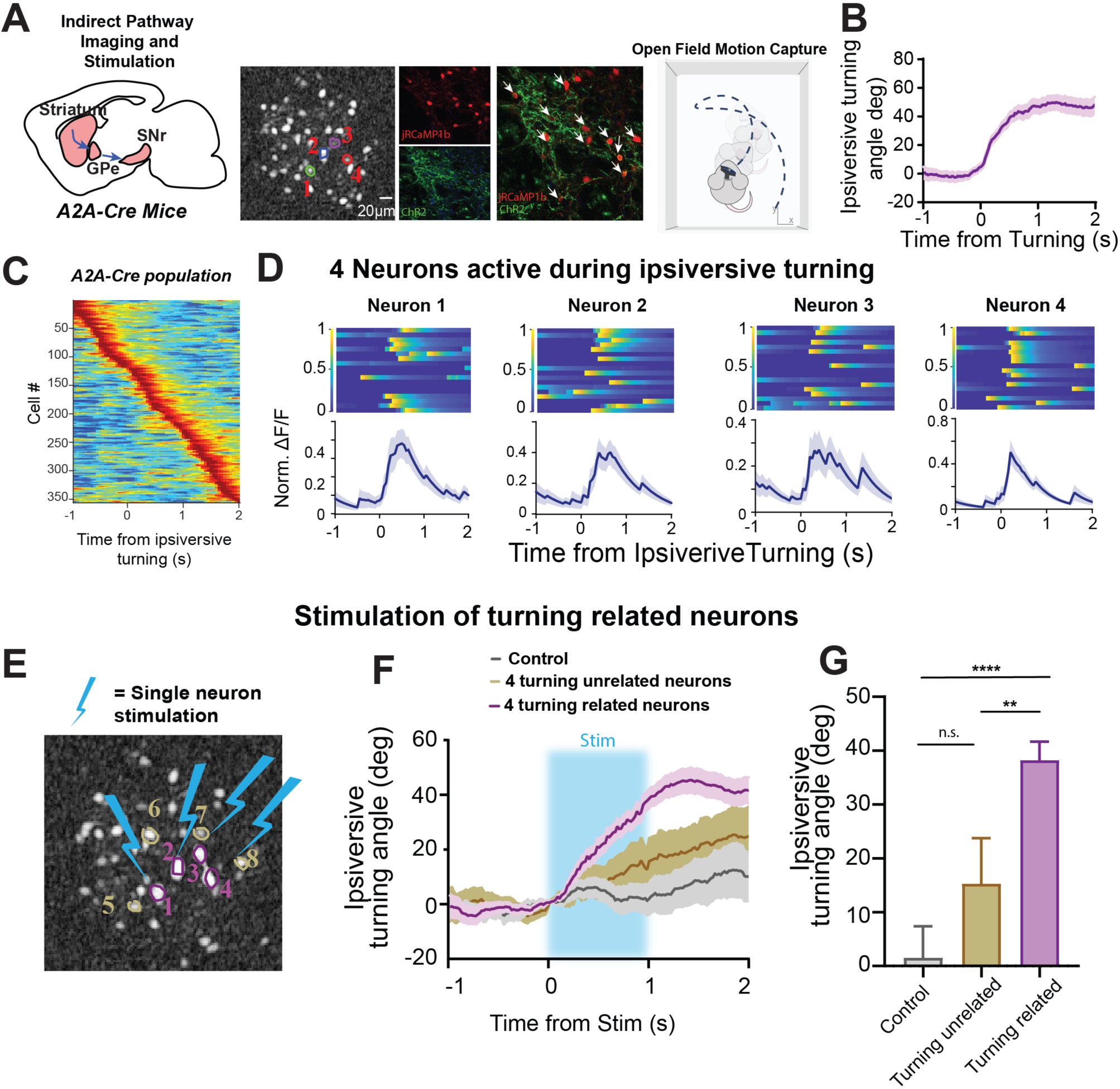
Selective stimulation of indirect pathway neurons that are active during ipsiversive turning recapulates ipsiversive turning (also see Video S2) **A)** 4 selected A2A+ DLS neurons from a representative mouse in an open field arena were stimulated. Neurons co-expressing both ChR2 and jRCaMP1b are shown in the histological images. **B)** Behavioral data was aligned to the onset of ipsiversive turning relative to the hemisphere being recorded from (3 A2A-cre mice, 5-28 trials per mouse). **C)** All recorded A2A+ neurons (3 A2A-cre mice, total of 354 neurons, >50 neurons per mouse) aligned to ipsiversive turning and sorted according to turning onset. **D)** 4 neurons from a representative mouse were selected that were active during ipsiversive turning. Data was aligned to ipsiversive turning onset. **E)** The same 4 neurons from one sample mouse were stimulated. Four neighboring neurons which not high related with turning were also stimulated. **F)** Optogenetic excitation of the 4 selected neurons produced significantly more ipsiversive turning compared to controls (no opsin expression) or unrelated neurons (3 A2A-cre mice, 6-30 trials per mouse for stimulated neurons, 3-16 trials per mouse for unrelated neurons; 2 control mice, 6-13 trials per mouse). **G)** Stimulation increased ipsiversive turning (one-way ANOVA: Stim, F(2,56) = 11, p<0.0001). Tukey’s post hoc analysis revealed stimulation of turning-related neurons produced greater ipsiversive turning compared to stimulation of unrelated neurons (p = 0.0096) or of controls (p <0.0001), but stimulating unrelated neurons did not produce more ipsiversive turning than controls (p=0.2932). All error bars indicate SEM.

Moreover, the effect of selective stimulation was also stable across time. We had selectively stimulated a specific ensemble of neurons on one day, and replicated the behavioral effect by stimulating the same neurons 40 days later (**Supplementary Figure 8**, 2 A2A-cre mice). There was no significant difference in the ipsiversive turning that was elicited on day 1 compared to day 40.

### Synthesizing sequences and sweeping patterns of stimulation

To test the effect of arbitrary stimulation patterns, we used two sequential patterns to activate turning-related neurons: either lateral to medial (LM, starting with the most lateral neuron) or medial to lateral (ML, starting with the most medial neuron) (**Figure 6B-C**). Both sequences produced contraversive turning (**Figure 6D**).

**Figure 6.**
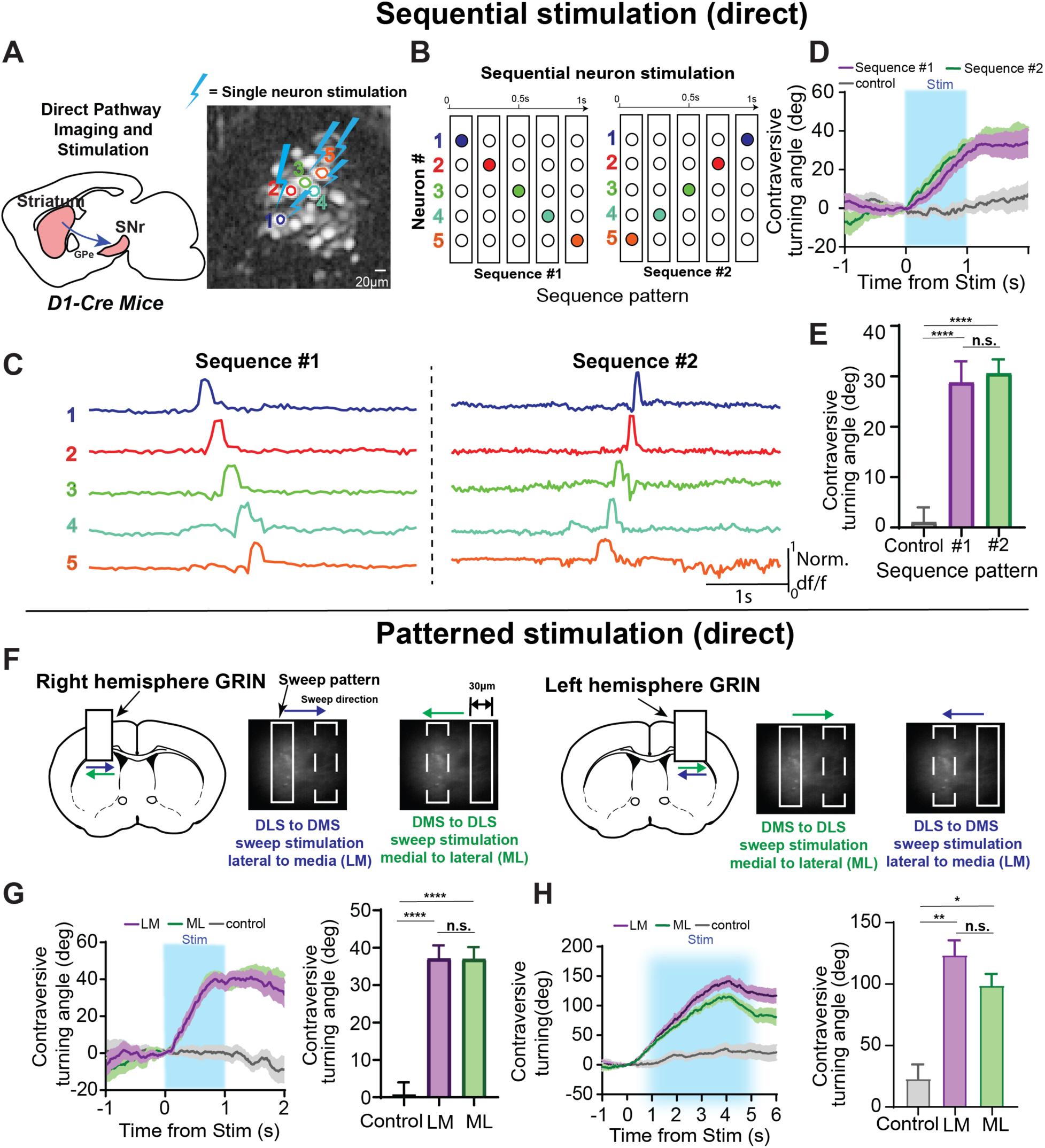
Sequential and patterned stimulation of the direct pathway causes contraversive turning. **A)** D1-Cre mice were used with MAPSI for single neuron sequence stimulation from a representative mouse. **B)** Five neurons were stimulated in two different sequences. The schematic shows the order and sequence of neurons that were stimulated. The two sequences either progressed laterally-to-medially (sequence #1) or medially-to-laterally (sequence #2) direction. **Also see Video S3. C)** Calcium traces for the selected neurons from each sequence, showing the time course of their activation. **D-E)** Stimulation of the 5 neurons in both sequences greatly increased maximum contraversive turning compared to controls, but there was no difference of the direction of the sequence (one-way ANOVA, F(2,60) = 4.112, p < 0.0001. Tukey’s post hoc analysis revealed that both sequences produced significantly more contraversive turning than controls (p =<0.0001, p < 0.0001), but were not different from each other (p = 0.9277) (3 D1-Cre mice, 5-25 trials for #1, 4-20 trials for #2, 2 control mice, 6-18 trials per mouse). **F)** Schematic of sweeping stimulation experiment where a 20 μm-wide bar moved horizontally, from lateral side to medial side (LM) or from medial side to lateral side (ML). **G)** Sweeping stimulation for 1 second increased contraversive turning compared to controls (left). There was a significant main effect of stimulation vs. non-stimulation (one-way ANOVA F(2,58) = 0.1825, p = 0.8336. Tukey’s post hoc analysis revealed that sequences LM vs ML was not significant: p =0.9996; but LM vs controls: p < 0.0001, and ML vs. controls, p < 0.0001, were significant; 3 D1-Cre, 4-17 trials LM, 6-25 trials for ML; 2 controls, 5-19 trials per mouse). **H)** Sweeping stimulation for 5 seconds significantly increased contraversive turning compared to controls. There was a main effect of stimulation (F_(2,12)_ = 5.88, p = 0.017). Tukey’s post hoc analysis however revealed no significant difference between sequence #1 and #2 (p = 0.74), but a significant difference between sequence #1 and controls (p = 0.0023) and between sequence #2 and controls (p = 0.049) (3 D1-Cre mice, LM: 5-21 trials per mouse, ML: 5-25 trials per mouse; 3 controls, LM and ML, 6-20 trials per mouse). All error bars indicate SEM.

MAPSI also makes it possible to produce arbitrary spatiotemporal patterns of light to sculpt neural activity. We programmed the DMD to produce rectangular sweeping patterns that covered ∼20 % of the FOV. We used two different directions (ML or LM) across the FOV in both hemispheres (**Figure 6F**). In D1-Cre mice (N=3) either pattern produced contraversive turning, but there was no significant difference between different sweep directions (ML or LM) regardless of hemisphere stimulated (**Figures 6G**,**6H**).

We next performed the same experiments using A2A-Cre mice (**Figure 7A**). We identified neurons that were active during ipsiversive turning, and used the same sequential patterns as used in figure 6 (**Figure 7B**). We also verified the calcium activity during the sequential pattern stimulation (**Figure 7C**). Both sequences produced ipsiversive turning with the LM sequence (#1) producing greater ipsiversive turning (**Figure 7D**). We then used the sweeping patterns used in **Figure 6**: ML or LM sweep across the FOV in both hemispheres (**Figure 7F**). We found that the rectangular sweep both ML and LM sweeps would significantly increase ipsiversive turning compared to controls, but there was no significant difference between different sweep directions (ML or LM) (**Figures 7G**). If we swept across the FOV over 5 seconds, both ML and LM sweeping stimulation produced more turning than controls, and LM produced significantly greater turning compared to ML sweeps (**Figure 7H**).

**Figure 7.**
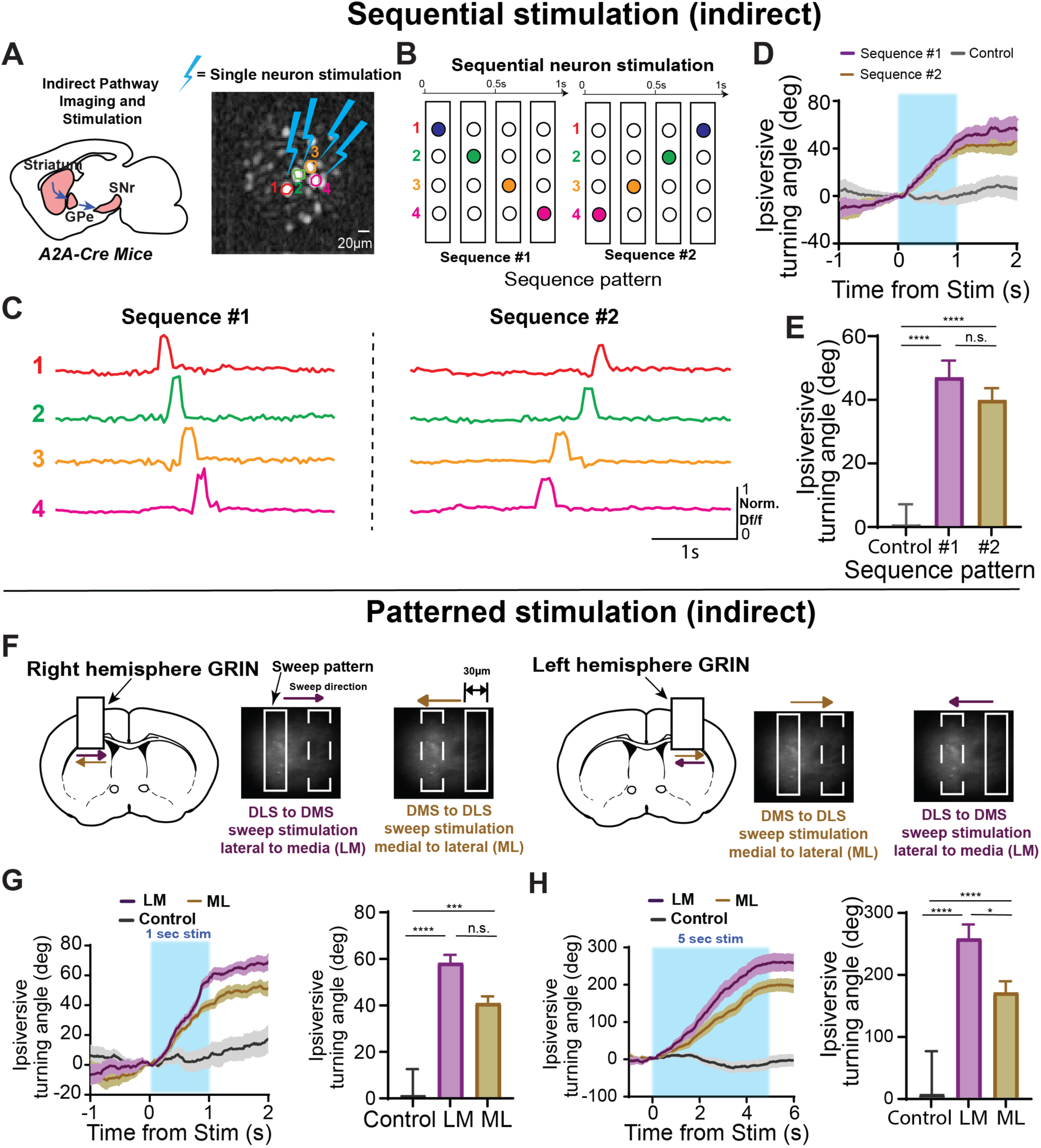
Sequential and patterned stimulation of the indirect pathway causes ipsiversive turning. **A)** A2A-Cre mice were used with MAPSI for single neuron sequence stimulation. **B)** 4 neurons were optogenetically excited in two different sequences. The schematic shows the order and sequence of neurons that were stimulated. Rows indicate individual neurons and columns indicate individual time points. The two sequences either progressed laterally-to-medially (sequence #1) or medially-to-laterally (sequence #2). **C)** Calcium traces for the 4 selected neurons from each sequence, showing the time course of their activation. **D-E)** Stimulation of the 4 neurons in both sequences significant increased ipsiversive turning compared to controls (one-way ANOVA, F_(2,51)_ = 0.63, p=0.5367, #1 vs #2: p = 0.6191, #1 vs control: p<0.0001, #2 vs control: p < 0.0001, 3 A2A-cre mice, 3-19 trials of LM per mouse, 3-17 trials of ML per mouse; 2 controls, 6-18 trials per mouse). **F)** Schematic of sweeping stimulation experiment. A 20 μm-wide bar moved horizontally, from lateral side to medial side (LM) or from medial side to lateral side (ML). Also see Video S4. **G)** Sweeping stimulation for 1 second significant increased ipsiversive turning compared to controls. Sweeping from lateral to medial also produced more ipsiversive turning. Ipsiversive turning was significantly higher than controls for both sequences, however were not significant compared to each other (one-way ANOVA, F_(2,12_) = 14.79, p= 0.0006. Tukey’s post hoc analysis reviewed that sequence #1 was not significantly different than sequence #2, p = 0.24), but sequence 1 (p = 0.0005 and sequence 2 (p = 0.0092) were significantly greater than controls 3 A2A-cre, 4-12 trials of LM per mouse, 3-12 trials of ML per mouse, 2 controls, 4-12 trials per mouse). Schematic representation of sweeping photo-stimulation. **H)** Sweeping stimulation for 5 seconds significantly increased ipsiversive turning compared to controls, and the LM sweep produced more ipsiversive turning compared to the ML sweep. (one way ANOVA, F_(2,12)_ = 47.68, p < 0.0001,3 A2A-cre mice, 3-10 trials per mouse for LM and 4-17 trials for ML; 2 control mice, 6-13 trials per mouse). Tukey’s post hoc analysis reveals that both sequences produced more ipsiversive turning than control (p < 0.0001 for both), LM vs ML: p=0.0132. All error bars indicate SEM.

## Discussion

The MAPSI system can 1) provide near-cellular resolution stimulation (**Figures 1-2**), 2) stimulate and record from selected neurons and simultaneously record activity in other neurons in the FOV **(Figure 3)**, 3) identify neurons active during a specific behavior and then selectively activate the relevant neuronal ensembles to reproduce the behavior (**Figures 4-5**), 4) recreate the recorded spatiotemporal pattern with stimulation, and 5) synthesize arbitrary stimulation patterns in the FOV (**Figures 6-7**).

A major challenge in optogenetics is to control the spatial location and extent of photo-stimulation. Traditionally light delivery is achieved with flat-faced optical fibers which illuminate a relatively fixed brain volume around the tip of the fiber. Although recently developed optogenetic methods can control the extent of light delivery, it is difficult to generate patterned stimulation with high resolution in freely behavior mice without using a 2P setup ^41^. Although 2P methods can achieve cell-specific stimulation, they require expensive setups and often cannot be performed in freely moving animals. In contrast, MAPSI makes it possible to synthesize complex spatiotemporal sequences of neural activity in any brain region in freely moving animals.

An important caveat is that single cell resolution has not been demonstrated in our system, since we cannot adjust the stimulation along the z-axis. As described above, a 10 µm spot on the selected neuron could potentially penetrate up to 30 µm below the surface of the GRIN lens. Given the size of SPNs (∼ 12-15 µm diameter), it is possible that additional neurons located just beneath the selected neurons were also activated. However, even in this scenario there cannot be many neurons affected. Depending on the size of the neuron and the density in the brain area in question, MAPSI can approach cellular resolution in many instances. While it is possible that, as opsin expression may not be confined to the soma, some opsin-expressing neighbor neurons might be excited by the photo-stimulation, this possibility would be less likely if soma-targeted opsins are used^42^.

By using a DMD, we were able to generate precise beamlets that can target single, or multiple, user-selected neurons with single-cell resolution. We tested this design in freely moving mice by simultaneously recording and stimulating direct and indirect pathway neurons in vivo. We were able to replay the recorded calcium activity in a small group of neurons, and reproduce the same behavior (**Figures 6-7**). This is the first demonstration that selective stimulation of a few SPNs could result in turning behaviors. In either direct or indirect pathways, activation of as few as 3 SPNs could produce significant turning, contraversive for dSPNs and ipsiversive for iSPNs. The amount of turning depends on the number of turning-related neurons activated, and activation of 5 neurons produced a comparable amount of turning as stimulation of the entire FOV (**Supplementary Figures 5 and 6**). This is not surprising since in the FOV the number of SPNs that are specifically related to turning is low, and stimulating the whole field would additionally activate neurons with presumably other functions. These results suggest that the number of SPNs involved in generating a specific action is much lower than expected.

MAPSI represents a major advance over existing systems for simultaneous imaging and photo-stimulation. Current systems that are capable of patterned stimulation and imaging usually require large microscopes and are difficult to use in freely-moving animals^13,43^. The best example of a 1P system is a fiberscope system developed by Szabo and colleagues^22^. But their system has several limitations. First, it requires a confocal microscope as well as two separate lasers, whereas our system does not require a separate microscope and uses only one laser. Secondly, with the fiberscope system, lateral and axial resolution can be degraded when the animal is moving with the optic fiber. Moreover, the wavelength for ChR2 stimulation overlaps with the excitation wavelength of the calcium indicator (GCaMP5), so some neurons could be excited during calcium imaging. The FOV for stimulation and recording using their fiberscope system is also limited by the properties of the coupled micro-objective. While the FOV of the fiberscope is similar to MAPSI (∼250 µm), our system uses a GRIN lens, which makes it possible to move the baseplate to move the FOV to a different region covered by the lens. Thus many neurons can be recorded and stimulated from a single animal. There is also a commercially available 1P fiberscope (OASIS Implant) based on the work by Szabo et al (https://www.mightexbio.com/products/oasis/oasis-implant/). This system has the advantage of a large FOV (0.5-3.2 mm) and a small implant (0.7 g), but has similar limitations as described above for the system from Szabo et al.

In MAPSI, because the dual-beam paths are independent of each other, neurons selected for stimulation can be simultaneously recorded along with the activity of other neurons in the FOV. MAPSI can be used as a tool in investigating mouse models of neurological and psychiatric disorders. For example, it would be useful to characterize pathological neural activity in disease models, to determine whether they play a causal role in the key symptoms, and also to rescue function by artificially introducing more normal patterns of neural activity. In short, given its small size, low cost, and portability, MAPSI provides a powerful new tool for interrogating neural circuit function in freely moving animals.

## Methods

### Experimental subjects

All experimental procedures were approved by the Animal Care and Use Committee at Duke University. Male D1-Cre mice (Jackson Labs: Drd^1tm2.1Stl^) and A2A-Cre (Adora2A^tm1Dyj/J^) mice were used. All mice were between 3-8 months old, group housed, and maintained on a 12:12 light cycle. Testing was always performed during the light phase.

### Viruses

pAAV.CAG.Flex.NES-jRCaMP1b.WPRE.SV40 was a gift from Douglas Kim & GENIE Project (Addgene viral prep # 100849; http://n2t.net/addgene:100849 ; RRID:Addgene 100849).. pAAV-EF1a-double floxed-hChR2(H134R)-EYFP-WPRE-HGHpA was a gift from Karl Deisseroth (Addgene plasmid # 20298 ; http://n2t.net/addgene:20298 ; RRID:Addgene_20298)

### Surgery and Histology

Mice were initially anesthetized with 5.0% isoflurane and maintained at 1-2% during surgery. A craniotomy was made to allow implantation of the GRIN lens (Bregma +0.0-1.0 mm AP, ±2.0-2.7 ML). Pulled pipettes were used to inject the virus using a Nanoject III injector (Drummond Scientific, USA). The first virus injection (250 nl of pAAV.CAG.Flex.NES-jRCaMP1b.WPRE.SV40) was injected at two sites (+0.25, +0.75 AP, 2.5 ML) each with 5 depths (2.8-2.0 DV). Injections were made at a rate of 1 nl per second. The second injection (250 nl of AAV(9)-EFIa-DIO-hChR2(H134R)-EYFP) was then injected at the same coordinates. The injection pipette was always left in place for three minutes after each injection to allow for maximum absorption before it was retracted.

After the virus injection, aspiration was performed from brain surface, and a GRIN lens (1.8 mm x 4.3 mm, Edmund Optics) was implanted in the DLS above the injection site. The lens was secured to the skull using dental cement and covered with Kwik-Sil to protect the lens surface. 5-6 weeks after the GRIN lens implantation, base plating was performed under visual guidance of the calcium signal to determine the best FOV.

After the completion of experiments, mice were transcardially perfused with 0.1M phosphate buffered saline (PBS) followed by 4% paraformaldehyde (PFA) to confirm placement and viral expression. The brains were then transferred to a 30 % sucrose solution and sliced coronally using a cryostat (Leica CM1850). Slices were mounted with DAPI-mounting medium (Vector Laboratories, Vectashield, cat. no. H-1800) to identify the nuclei of neurons. Slices were then imaged using an inverted confocal microscope (Zeiss LSM780 and LSM880) for zoomed in images or an upright epifluorescence microscope for whole brain images (Axio Imager.M1 - Zeiss).

### Filters and LED Control

A lime LED and a blue laser are both used in MAPSI. A LUXEON Rebel Color lime LED (LXML-PX02-0000) filtered with a 540-580nm excitation LED filter (Chorma ET560/40x) is used for calcium imaging, while a 473 nm blue laser filtered with an excitation 450-490nm laser filter (Chroma ET470/40x) is used for optogenetic stimulation. The two colors are combined through the excitation dichroic mirror (Chroma 59003bs) by merging them in the excitation light path. The main dichroic mirror (Chroma 69013bs) reflects the excitation light into the GRIN lens, while the fluorescence image passes through the emission filter. The emission filter (Chroma ET630/75) is designed to avoid crosstalk from the excitation light. In addition, an absorber (Chroma ET775/50x) is located beneath the DMD to avoid reflected blue light during the off-state that could potentially scatter into the excitation light path (**Supplementary Figure 2**).

The output from the lime LED is controlled by two parallel current chips (LT3092ETS8), both of which supply 2.9V of voltage and enough current to power the LED. The constant current source can provide up to 200 mA of current (up to 12 mW/mm^2^ measured beneath the GRIN lens) with a step size of 4 mA. A programmable potentiometer (MCP4018) is used to adjust the current using control signals from an Arduino UNO (Arduino), which communicates with the computer. Custom scripts are used to adjust the parameters of calcium imaging.

### Digital Micromirror Device (DMD) design and installation

For patterned stimulation, we used a DMD (Texas Instruments; DLP2010), which is a digitally controlled micro-opto-electromechanical system that modulates spatial light to generate different light stimulation patterns. This DMD has on its surface more than 400,000 microscopic mirrors arranged in a rectangular array, with a resolution ratio of 854 × 480 pixels. Each mirror has a hinge and hook beneath it, allowing it to rotate ±17° (relative to the flat surface), in order to switch between on- and off-states. The “on-state” is defined by each of the mirrors’ positions corresponding to the tilt angle θ (±17°), such that the reflected light is directed toward the excitation light path. When a mirror is positioned in the opposite direction, the mirror is said to be in the “off-state”. In the on-state, light from the blue laser is reflected into the lens. In the off state, light is reflected into the absorber rather than into the GRIN lens, preventing stimulation in the specified FOV (**Supplementary Figure 2**). The rotation angle should be limited, so that in the off-state, all the light will be reflected to the absorber without any light scattering. Therefore, the DMD rotation angle must be limited as follows:

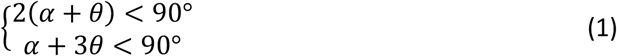

Where *θ* is the DMD rotation angle and *α* is the mirror tilt angle. We set *α* = 13° as the optimal rotation angle.

### Miniscope housing design and simulation in software

The miniscope body size is small, while the distance between the laser source and the GRIN lens is 20 mm. To ensure that we can generate precise patterns beneath the GRIN lens, and to have sufficient energy for excitation, the beam divergence for the laser light must be less than 4°. To generate a near perfect collimated light with sufficient energy in the small miniscope body, a laser fiber-head 100 μm in diameter is placed next to the focal point in order to gather all the energy and collimate the beam (**Supplementary Figure 3**). The collimation deviation is calculated by the following equation:

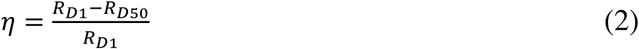

Where R_D1_ is the radius of the illumination field when the target screen is 1 mm from the outer surface of the collimation lens, R_D50_ is the radius of the illumination field when the target screen is 50 mm from the surface, and η is the collimation deviation. The smaller the deviation, the more precisely collimated the light is.

After the placement and operation of all the filters and lenses were confirmed, a 3D model housing all the components was designed in Solidworks software. To verify the optic simulation designs, we imported all the 3D models into the software program Tracepro (**Supplementary Figure 2**). The miniscope body was then printed using a 3D printer (Multi-Jet Fusion technology) with the material Nylon PA12.

### Miniscope control system design

MAPSI is based on the original UCLA open source Miniscope^23^. It includes: A CMOS imaging sensor, a PCB board, all the optical components such as the emission and excitation filters, the GRIN and achromatic lenses, and DAQ hardware and software. The image sensor resolution is 752px*480px and the frame rate is up to 60 Hz. However, the original Miniscope has been significantly modified to allow separate light paths. The CMOS, DAQ, and software are used after minor modifications.

The whole system contains an LED power source, a recording subsystem and a pattern control subsystem (**Supplementary Figure 1**). Two computers are needed: one that connects with all of the subsystems to send and receive data, as well as to send commands. A second computer is used to project the photo-stimulation patterns into the pattern control subsystem (**Supplementary Figure 2**). The first computer runs two programs: the original miniscope recording software, and the integration software application that controls the LED power subsystem, which sends synchronous commands to all of the other subsystems to ensure timestamps are all aligned. An Arduino UNO that is connected with this computer sends and receives the synchronous commands from/to the CMOS to control the LED current and the pattern. A National Instruments (NI) box is also connected with this computer, and sends TTL signal to control the laser generator (RL639T8-500).

The DMD was controlled by a custom designed driver board with a 60 cm enameled wire cable. The DMD driver board is fixed on the following-up turning holder with the HDMI interface board. One computer sent image data to the HDMI interface using a wireless HDMI Kit (Diamond VS75). The pattern control subsystem receives the pattern image data from the computer, and controls the DMD (Texas Instruments; DLPC3430 and DLPA2000). An HDMI data interface board transmits signals from the computer to the DLP3430.

### Determining z-axis resolution of MAPSI

The MAPSI z-axis resolution was determined in brain slices placed on a confocal microscope (Zeiss, LSM 880) equipped with a 20x (NA 0.8) objective. Using a vibratome, we cut 200 μm coronal sections from a mouse brain co-expressing RCaMP1b and ChR2. The slice was adhered to the bottom of a culture dish, and the GRIN lens was placed just above the slice. The GRIN lens and the tissue were then embedded together with 4% agarose gel. MAPSI was then fixed on the top of the GRIN lens, and attached to a stereotaxic frame to obtain a good focal pattern image (Figure 2B). A confocal microscope was used to image the slice from bottom of the dish while simultaneously using MAPSI to target single neurons from the top of the dish. While MAPSI is generating a pattern on the tissue, RCaMP1b images were acquired using 561 nm excitation laser and 2 detectors (1 PMT for the pattern and 1 GaAsP detector for RCaMP) (pixel size, 0.21 µl x 0.21 µl x 0.70 µl; pinhole size, 1 airy unit). The total energy detected beneath the GRIN lens was measured using optical power meter (PM100A).

### Calibration of MAPSI

To match DMD and CMOS pixel location, we used a 60 µm brain slice co-expressing jRCaMP1b and ChR2. GRIN lens was placed on the brain slice, and MAPSI fixed on the top of GRIN lens. Once an image with white spot (diameter = 10 pixels) and black background was projected on DMD, a spot fluorescence image was recorded on CMOS (**Supplementary Figure 4D**). As the white spot was moved in one axis, the movement of the fluorescence spot was also captured. Based on the DMD pixels (854*480) and the CMOS pixels (752*480), we were able to calibrate the white spot position and fluorescence spot position with the linear relationship:

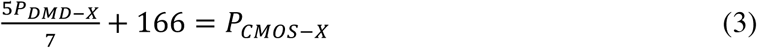

Where P_DMD-X_ is the x axis position on DMD, and P_CMOS-X_ is the x axis pixel position on CMOS.

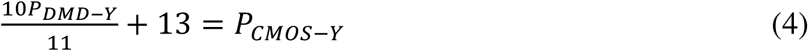

Where P_DMD-Y_ is the y axis position on DMD, and P_CMOS-Y_ is the y axis pixel position on CMOS.

Once a neuron is identified for stimulation, the pixels’ location of that neuron can be imported into equation (3) and (4) to calculate the matching pixels’ location on DMD. The projection image is then created with the calculated pixels.

### Open-field behavior recording and analysis

Mice were tested in an open field arena (25 cm x 25 cm). A high-speed camera (FLIR BFS-U3-04S2M) was placed above the platform to record the behavior of the mice at 50 frames/s. The DMD driver and HDMI interface board were fixed on the tuning holder, connected to the DMD using a 60c m enamelled wire cable which prevents tangling of the wires and allows the mice to move freely. Other wires (coaxial cable for CMOS, power wire for LED) pass through the center hole of ball bearing (**Supplementary Figure 4A**).

Behavioral data recorded by the cameras was analyzed using DeepLabCut ^40,44,45^. After labeling several frames (skeleton with markers on the head and back of the mice), the behavior data can be calculated by transfer learning with deep neural networks. In DeepLabCut, 3 markers (head, body, and tail) were labeled in each open-field behavioral movie. 200 samples of frames were auto selected from each video and 18 videos was taken for training with the 200,000 training iterations. The test error was 4.37 pixels and the training error was 1.63 pixels. The resulting data was then imported into MATLAB where behavioral variables, such as displacement and turning angle, were created using a custom script. The processed data was then imported into Neuroexplorer 5 along with the calcium imaging data for further analysis

### Statistical Analyses

All statistical analyses were performed in GraphPad Prism 7.0. All error bars represent standard error of the means (SEM). Significance levels were set to P < 0.05. Significance for comparisons: *P < 0.05; **P < 0.01; ***P < 0.001; ****P < 0.0001.

### Calcium imaging acquisition and analysis

A CMOS imaging sensor, a data acquisition system (DAQ), and a USB host controller were used for calcium imaging. Images were acquired at 20 frames per second and recorded to uncompressed “.avi” files using the DAQ software. The videos were then imported into MATLAB for non-rigid motion correction ^45^, followed by deconvolution with the constrained non-negative matrix factorization (CNMF-E) algorithm ^46^. Seed pixels were initialized with a minimum local correlation value of 0.8 and a minimum peak-to-noise ratio of 10. The minimum number of nonzero pixels for each neuron was set at 10 based on the resolution of the CMOS sensor (1 µm/pixel). Both deconvolved and raw calcium traces were saved.

## Data and code availability

All data and Matlab codes used in the present study are available upon request.

## Ethical compliance

All research reported in this study was approved by Duke University Institutional Animal Care and Use Committee (protocol A254-19-11).

## Acknowledgements

This research was supported by NIH grant MH112883 to H.H.Y. We would like to thank Nicole Calakos, Shawn Je, and Mark Rossi for helpful comments on the manuscript.

## Author Contributions

H. H. Y. conceived the concept of all optical 1-photon system for freely moving mice. J.Z., N.K. and H.H.Y. designed the experiments. J.Z. designed, assembled, and calibrated the circuit board and MAPSI. J.Z and I.P.F performed testing in mice. J.Z, N.K., and K.B. analyzed data. N.K., I.P.F., K.B. and R.N.H performed surgeries. R.N.H & J.K. performed histology and confocal imaging. R.N.H., J.Z., & H.H.Y. wrote the manuscript.

