## Supplementary information for "All-optical imaging and patterned stimulation with a one-photon endoscope"


**Authors/Affiliations:**

Jinyong Zhang[1], Ryan N. Hughes[1], Namsoo Kim[1], Isabella P. Fallon[2], Konstantin Bakhurin[1], Jiwon Kim [1], Francesco Paolo Ulloa Severino[1,3], Henry H. Yin[1,2]†

**Affiliations:**

1. Department of Psychology and Neuroscience, Duke University
2. Department of Neurobiology, Duke University School of Medicine
3. Department of Cell Biology, Duke University School of Medicine

**Video S1**: Stimulation of a selected neuron in dorsal striatum.

**Video S2**: Stimulation of 4 indirect pathway neurons that are normally active during ipsiversive turning.

**Video S3**: Sequential stimulation of 5 direct pathway neurons that are normally active during contraversive turning.

**Video S4**: Sweeping stimulation of indirect pathway neurons.

### Supplementary Figures

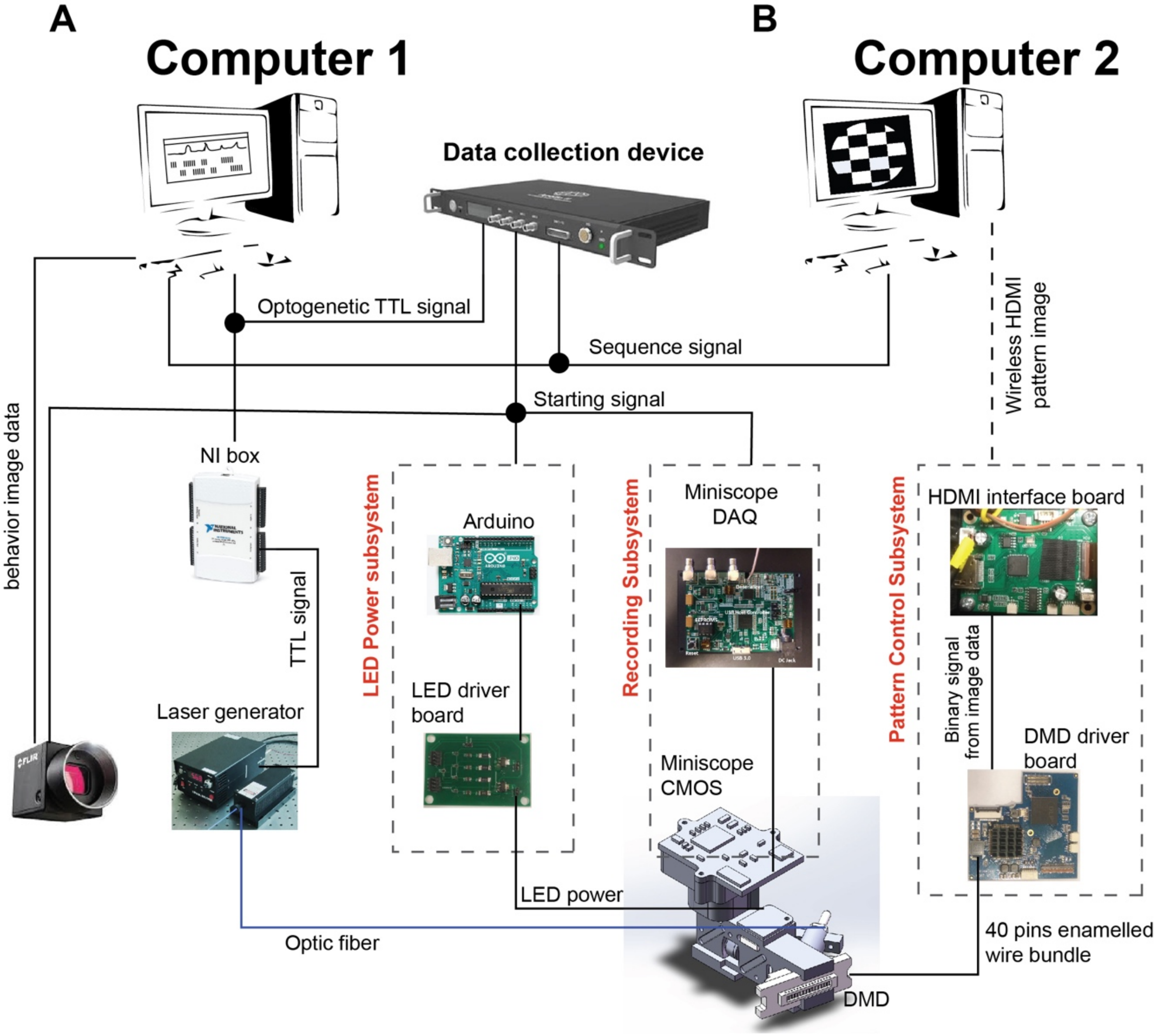


**Supplementary Figure 1. MAPSI design**

**A)** Computer 1 is primarily responsible for collecting data from the camera and MAPSI. It also commands all the subsystems to send and receive data. **B)** The second computer is primarily responsible for the pattern control subsystem, which projects pattern images onto the DMD.

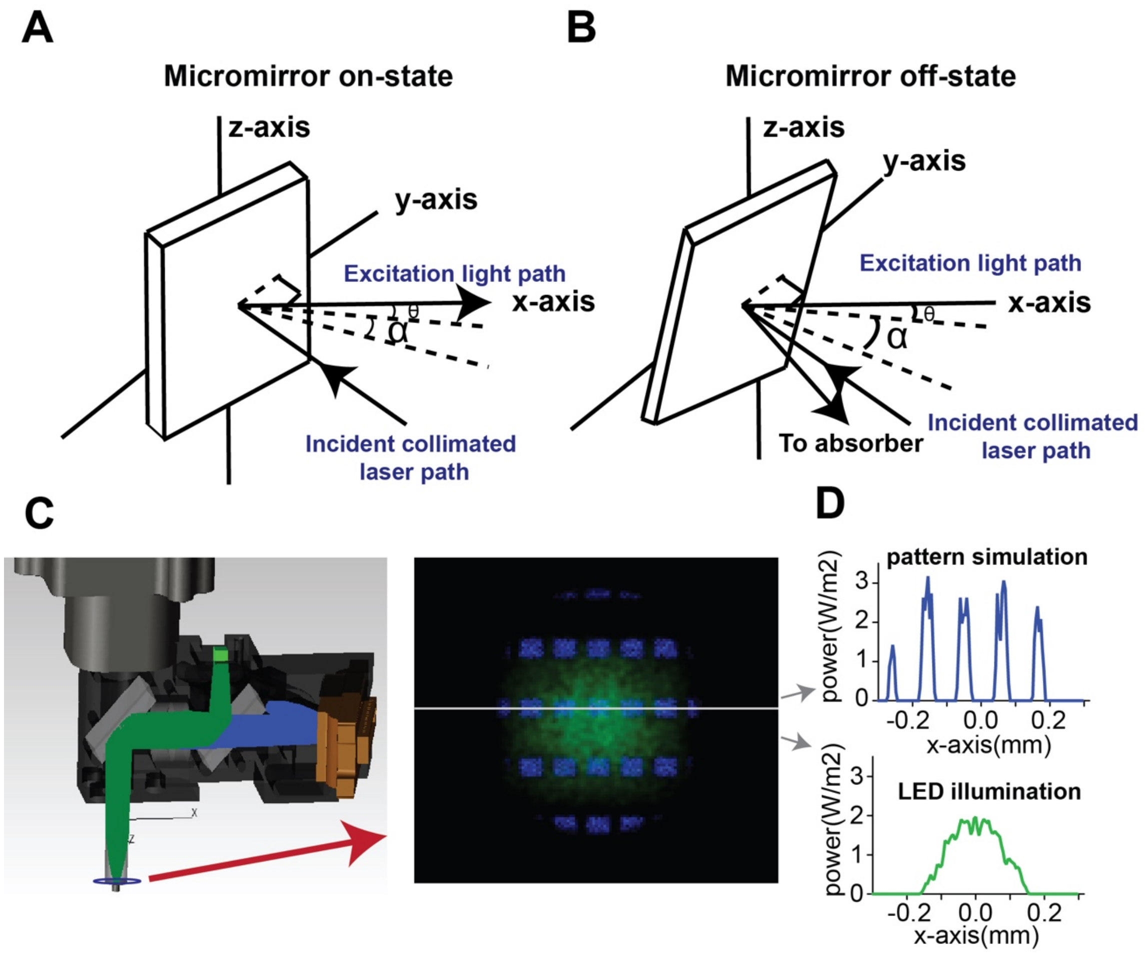


**Extended Figure 2. Configuration of the digital micromirror device (DMD)**

**A-B)** The DMD operates in two modes, on-state **(A)** and off-state **(B)**. In the on-state, the micromirrors are tilted at α (+17°). This allows the collimated light to be reflected to the excitation path. In the off-state, the mirrors in the DMD are tilted at -α(-17°), which causes the collimated light to be reflected towards the absorber rather than to the excitation path. The whole DMD must be rotated by θ (13°) to create enough room for the collimated lens. **C)** A model of the MAPSI pattern created in Tracepro, showing the pattern illumination produced by MAPSI. **D)** Simulation result from **C).** Light scattering is less than 5 μm. Peak power for each square is higher than 2 mW, which is above the threshold for exciting neurons.

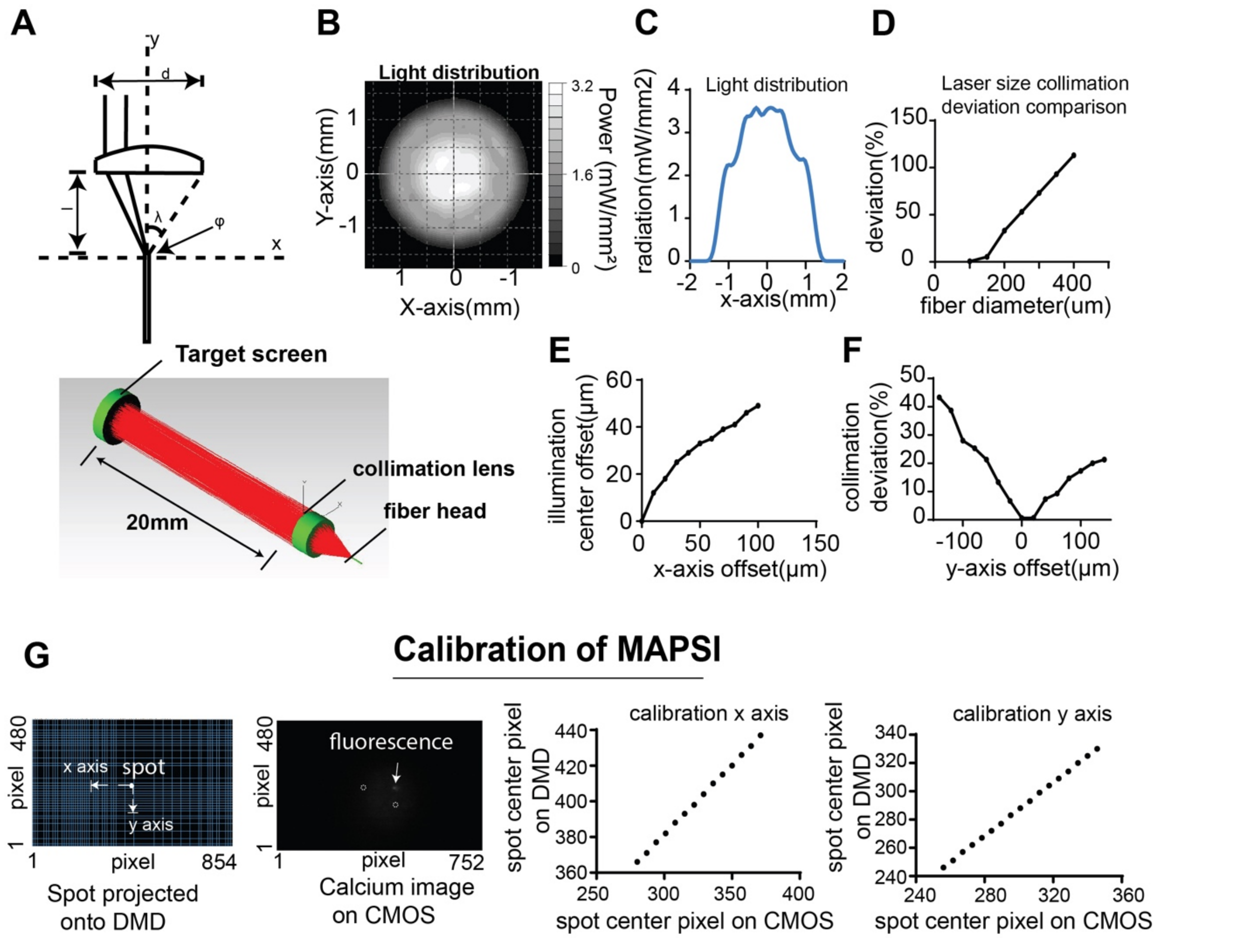


**Supplementary Figure 3. Laser collimation and calibration**

**A)** *Top)* Schematic drawing of laser light collimation. λ (20°) is radiation angle of the light from the fiber head. φ(100µm) is diameter of laser head. λ is measured by the power meter. The focal distance of the collimation lens is 3.5 mm, and *d* is the diameter of the lens. *Bottom*) Simulation settings and test of collimation using the software Tracepro. **B)** Energy distribution of the MAPSI. The peak power radiated on the target screen is 3.8 $mW/mm^2$, higher than what is needed to excite neurons. **D)** Simulation shows collimation deviation as a function of fiber diameter. A lower deviation is desirable as it results in higher resolution. **E)** Figure shows that x-axis offset increases the offset of illumination. Ideally, the fiber head should be positioned at the focal point of the collimation lens. **F)** As the offset increases, the collimation deviation also increases, reducing the precision of the stimulation. **G)** Calibration of MAPSI. An image with 10 pixels diameter white spot and black background was projected on DMD. While moving the spot, the fluorescence signal was recorded and used for calibration.

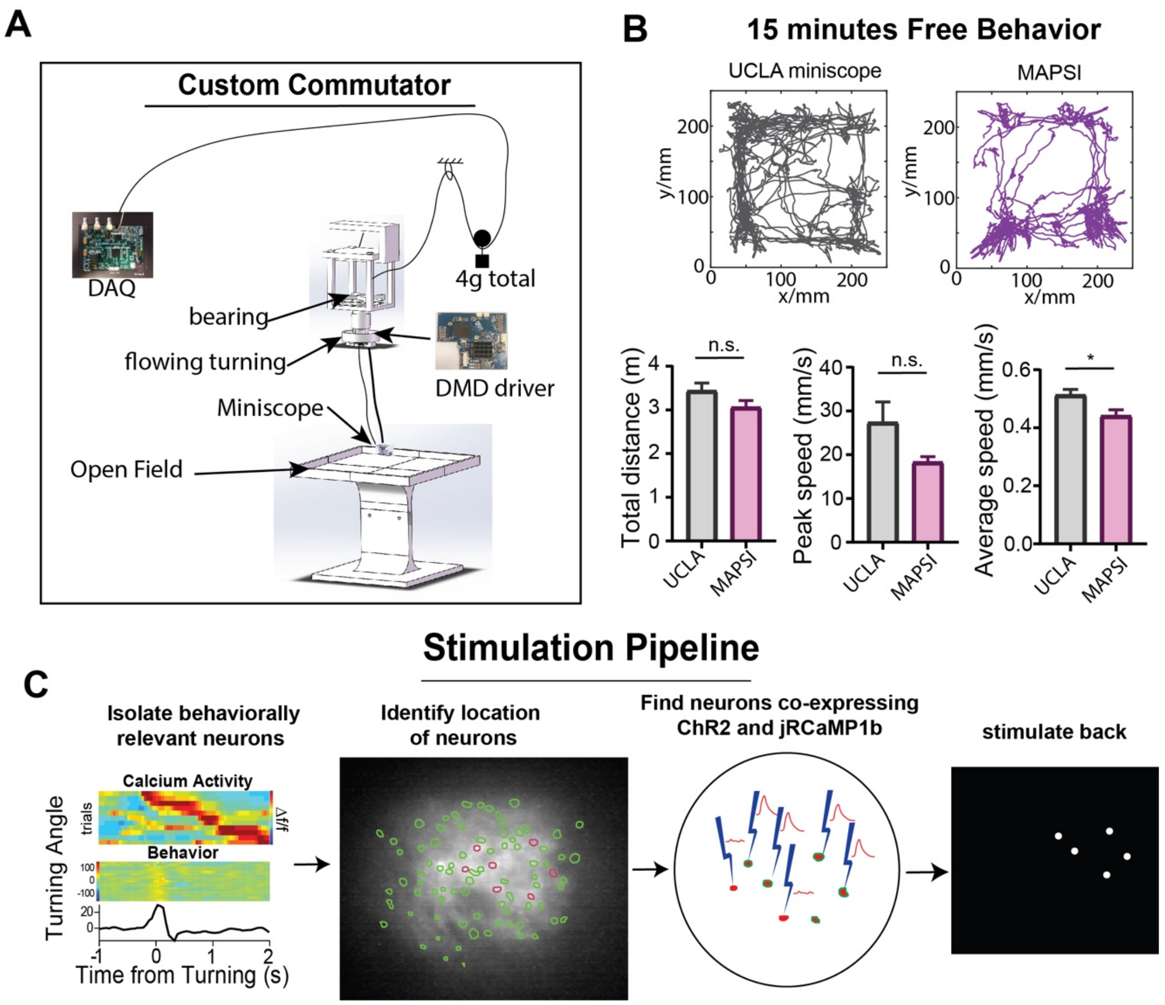


**Supplementary Figure 4. Free movement using MAPSI and stimulation pipeline.**

To reduce the weight of MAPSI carried by the mouse, a custom commutator was built. **A)** The commutator is attached to a 4g pulley, and allows the wires to rotate without being tangled. **B)** *Top*: Open field trajectory over 15 min of a representative mouse with 4 g UCLA miniscope compared to the trajectory of the same mouse carrying 7.8 g MAPSI. *Bottom*: Total movement distance, peak speed, and average speed (N=7, 4 D1-cre mice and 3 A2A-cre mice, 15 minutes in the same open field platform). Unpaired t test analysis revealed no significant difference between mice carrying UCLA Miniscope and MAPSI in distance (p=0.1274) or peak speed (p=0.0826), but mice carrying MAPSI showed reduced average speed (p=0.0238). * $p < 0.05$. **C)** Stimulation pipeline. Simultaneously record and analyze the calcium activity as well as the behavior. Isolate the behaviorally relevant neurons. Find the neurons that co-express both ChR2 and jRCaMP1b. Target neurons with co-expression and replay activity.

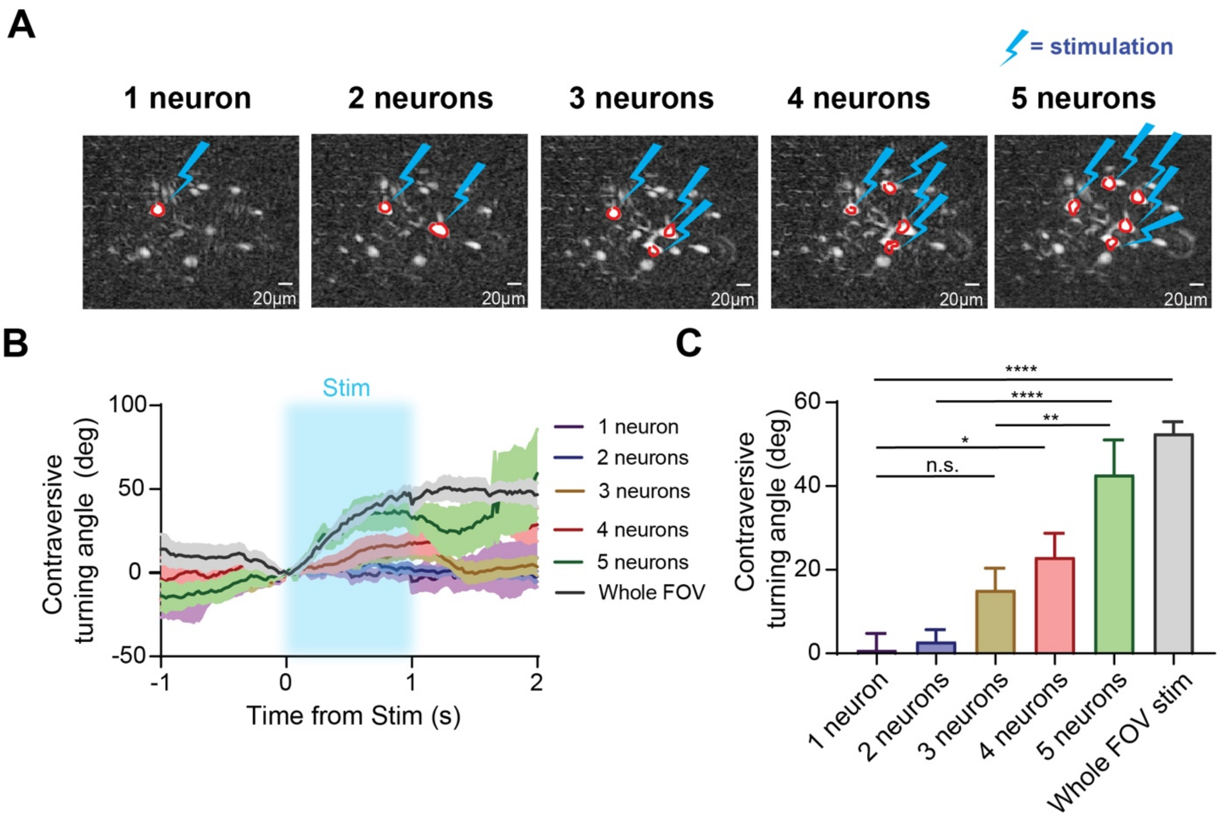


**Supplementary Figure 5. Contraversive turning behavior can be reliably elicited by stimulating as few as 3 dSPNs.**

**A)** Calcium imaging of the selected direct pathway neurons for photostimulation. Anywhere from one to five neurons were selectively stimulated. **B)** Optogenetically stimulating 3 or more neurons reliably produced contraversive turning. **C)** Turning was significantly higher when exciting 3 or more neurons (One-way ANOVA, $F_{(2,174)} = 15.39$, p = 0.0006). Tukey's *post hoc* analysis revealed no significant difference between 2 neurons and 1 neuron (p=0.9997), no significant difference between 3 neurons and 1 neuron(p=0.3750), significant difference between 4 neurons and 1 neuron (p=0.0454), significant difference between 5 neurons and 1 neuron (p<0.0001), and significant difference between whole FOV stim and 1 neuron (p<0.0001). (N= *2 D1-Cre* mice, 15 trials for 1 neuron, 13 trials for 2 neurons, 13 trials for 3 neurons, 11 trials for 4 neurons, 19 trials for 5 neurons, and 14 trials for whole FOV). * p < 0.05, ** p <0.01, *** p < 0.001, **** p < 0.0001.

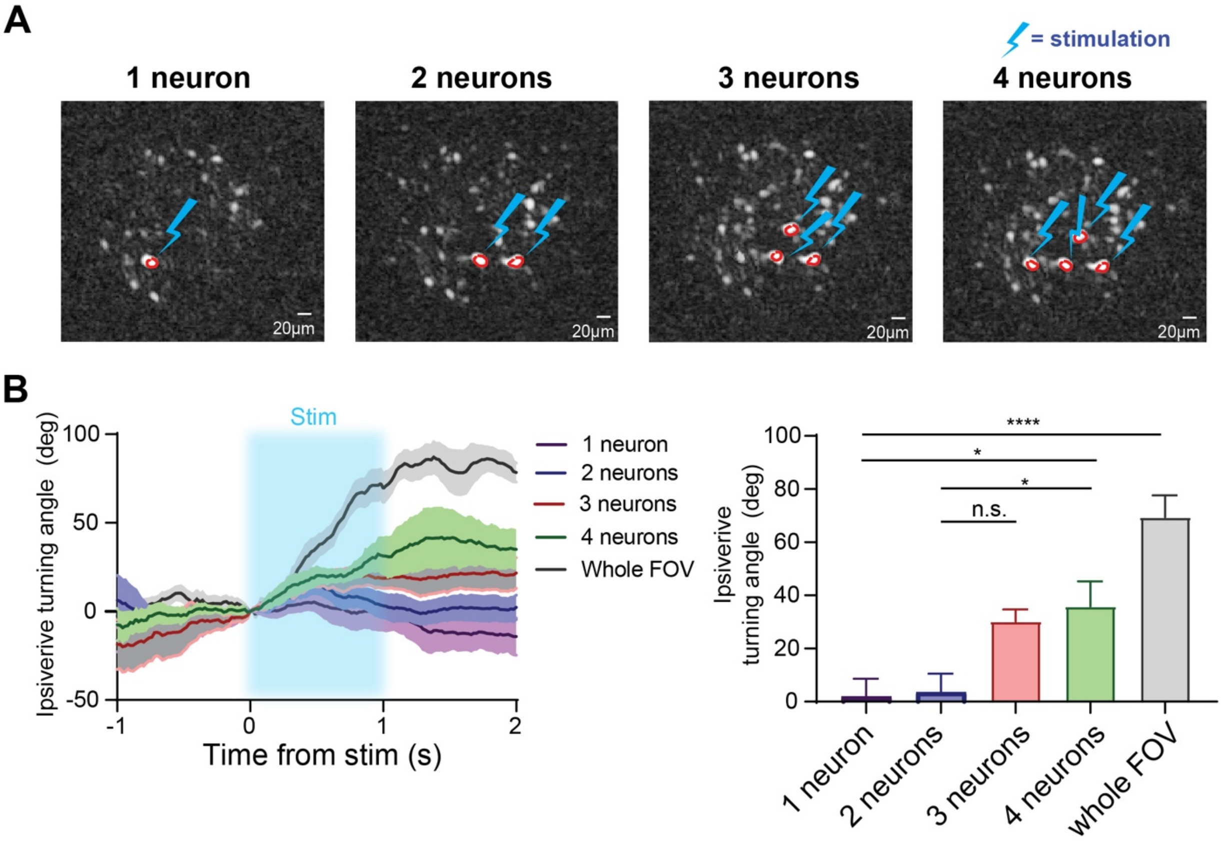


**Supplementary Figure 6. Ipsiversive turning behavior can be reliably elicited by stimulating as few as 3 iSPNs.**

**A)** Images of the selected indirect pathway neurons for photostimulation. **B)** Stimulating 3 or more neurons reliably produced contraversive turning. **C)** Higher number of stimulated neurons increased maximum ipsiversive turning angle (One-way ANOVA, $F_{(4,21)} = 6.104$, $p = 0.0037$). Tukey's *post hoc* analysis revealed no significant difference between 2 neurons and 1 neuron (p=0.9999), no significant difference between 3 neurons and 1 neuron(p=0.0953), significant difference between 4 neurons and 1 neuron (p=0.0303), and significant difference between whole FOV stim and 1 neuron (p<0.0001). 2 *A2A-Cre* mice, 10 trials for 1 neuron, 7 trials for 2 neurons, 9 trials for 3 neurons, 13 trials for 4 neurons, and 11 trials for whole FOV.

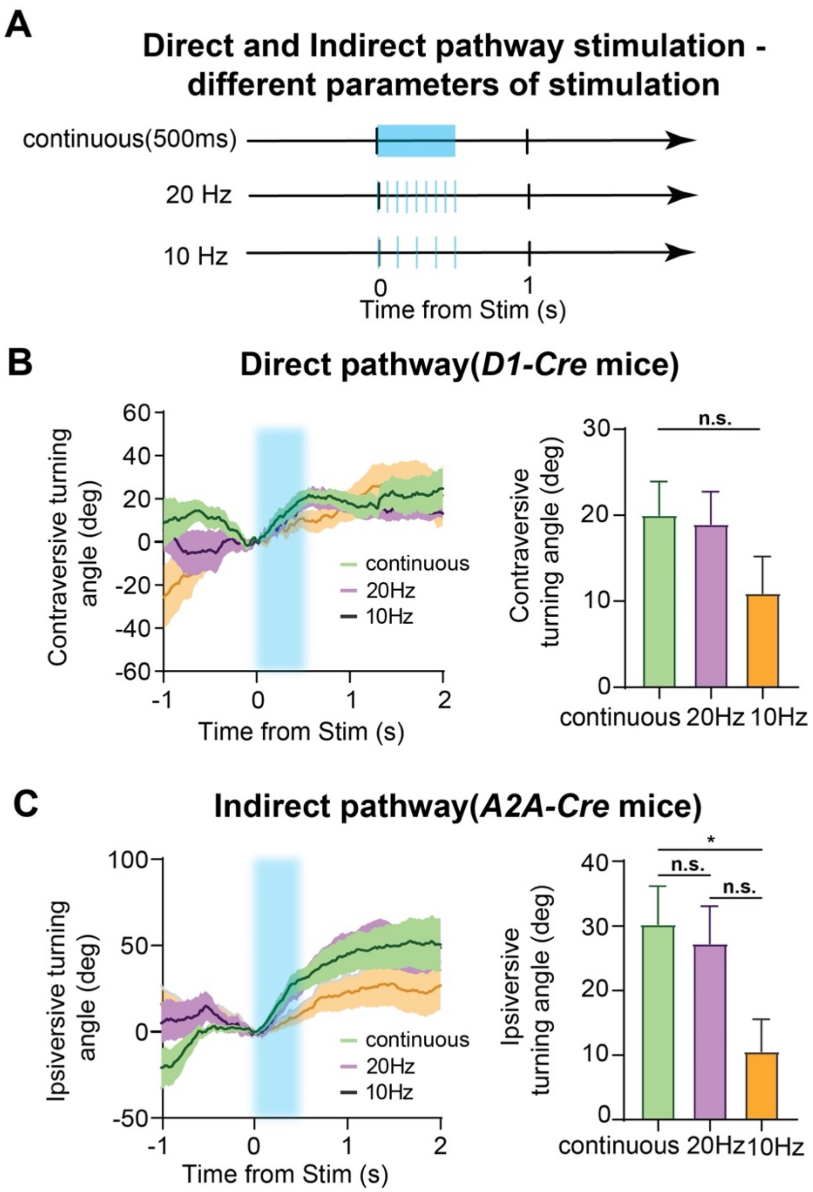


**Supplementary Figure 7. There is no significant difference in turning from direct and indirect pathway stimulation using different parameters of stimulation.**

**A)** We optogenetically stimulated neurons in the calcium imaging FOV with 3 different parameters: continuous 500ms, 20Hz (10 pulses in 500 ms), and 10 Hz (5 pulses in 500 ms). **B)** *Left)* Optogenetic excitation of direct pathway neurons in the DLS using the MAPSI significantly increased contraversive turning. *Right)* There was no significant stimulation effect (One-way ANOVA, $F_{(2,24)}$=1.315, $p$ = 0.2872) (N=2 *D1-cre* mice, 10 trials for 10Hz, 8 trials for 20Hz, and 12 trials for continuous). **C)** *Left)* Optogenetic excitation of indirect pathway neurons in the DLS using the MAPSI significantly increased ipsiversive turning behavior (One-way ANOVA, $F_{(2,24)}$=4.099, $p$= 0.0294; continuous vs 20Hz, $p$= 0.9367; continuous vs 10Hz, $p$=0.0407, 20Hz vs 10Hz, $p$=0.1080) (2 *A2A-cre* mice, 12 trials for 10Hz, 8 trials for 20Hz, and 8 trials for continuous). * $p < 0.05$.

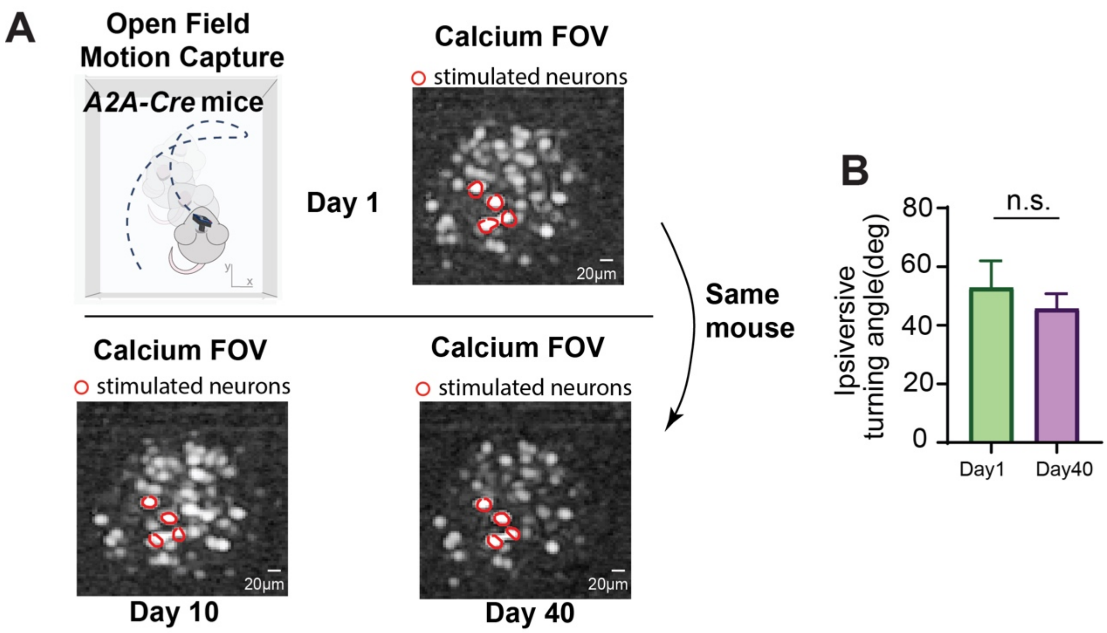


**Supplementary Figure 8. Behavioral effect of stimulation is stable across time.**

**A)** *Top*, a representative A2A-cre mouse was placed in an open field arena where several neurons were selected for stimulation on day 1 of a simultaneous stimulation and recording experiment. Shown on the right are the neurons selected for stimulation. *Bottom*, the same mouse was then tested 10 days and 40 days later in the open field arena, and the same neurons from day 1 were selectively stimulated. **B)** There was no significant difference in turning from day 1 compared to day 40 ($t_{(14)} = 0.71$, $p = 0.49$, 2 *A2A-cre* mice, 12 trials on day 1, 13 trials on day 40)

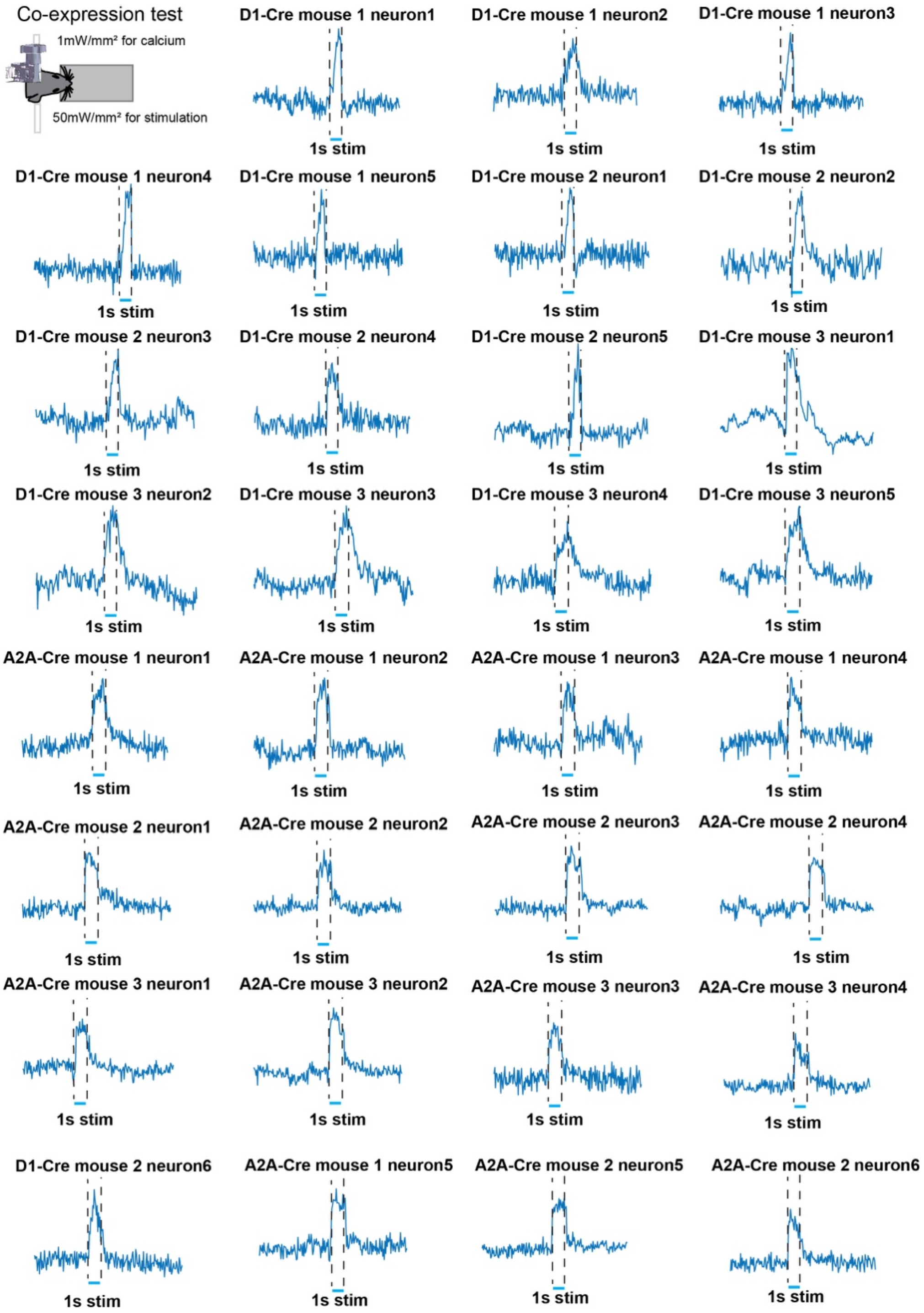


**Supplementary Figure 9. Traces from all neurons co-expressing jRCaMP1b and ChR2 (see Figure 2).**

Photostimulation of neurons that co-expressed jRCaMP1b and ChR2 when the mouse was anesthetized. Excitation power for imaging is ~1 mW/mm2, and stimulation power is ~50 mW/mm2 using 10 μm diameter laser spot. 3 D1-Cre mice, 3 A2A-Cre mice. N= 31 neurons.
